# TREE-QMC: Improving quartet graph construction for scalable and accurate species tree estimation from gene trees

**DOI:** 10.1101/2022.06.25.497608

**Authors:** Yunheng Han, Erin K. Molloy

**Affiliations:** Department of Computer Science, University of Maryland, College Park, 20742, USA; University of Maryland Institute for Advanced Computer Studies, College Park, MD 20740

## Abstract

Summary methods are one of the dominant approaches for estimating species trees from genome-scale data. However, they can fail to produce accurate species trees when the input gene trees are highly discordant due to gene tree estimation error as well as biological processes, like incomplete lineage sorting. Here, we introduce a new summary method TREE-QMC that offers improved accuracy and scalability under these challenging scenarios. TREE-QMC builds upon the algorithmic framework of QMC (Snir and Rao 2010) and its weighted version wQMC (Avni et al. 2014). Their approach takes weighted quartets (four-leaf trees) as input and builds a species tree in a divide-and-conquer fashion, at each step constructing a graph and seeking its max cut. We improve upon this methodology in two ways. First, we address scalability by providing an algorithm to construct the graph directly from the input gene trees. By skipping the quartet weighting step, TREE-QMC has a time complexity of *O*(*n*^3^*k*) with some assumptions on subproblem sizes, where *n* is the number of species and *k* is the number of gene trees. Second, we address accuracy by normalizing the quartet weights to account for “artificial taxa,” which are introduced during the divide phase so that solutions on subproblems can be combined during the conquer phase. Together, these contributions enable TREE-QMC to outperform the leading methods (ASTRAL-III, FASTRAL, wQFM) in an extensive simulation study. We also present the application of these methods to an avian phylogenomics data set.

## Introduction

Estimating the evolutionary history for a collection of species is a fundamental problem in evolutionary biology. Increasingly, species trees are estimated from multi-locus data sets, with molecular sequences partitioned into (recombination-free) regions of the genome (referred to as loci or genes). A popular approach to species tree estimation involves concatenating the alignments for individual loci together and then estimating a phylogeny under some model of molecular sequence evolution, like the Generalized Time Reversible (GTR) model (Tavaré 1986).

Standard models assume the genes have a shared evolutionary history; however, this is not necessarily the case. The evolutionary histories of individual genes (referred to as gene trees) can differ from each other due to biological processes (Maddison 1997). Incomplete lineage sorting (ILS), one of the most well-studied sources of gene tree discordance, is an outcome of genes evolving within populations of individuals, as modeled by the multi-species coalescent (MSC) (Pamilo and Nei 1988; Rosenberg 2002; Degnan and Salter 2005). Concatenation-based approaches to species tree estimation can be statistically inconsistent under the MSC (Roch and Steel 2015). Moreover, simulation studies have shown concatenation can perform poorly when the amount of ILS is high (e.g. Kubatko and Degnan 2007). ILS is expected to impact many major groups, including birds (Jarvis et al. 2014), placental mammals (McCormack et al. 2012), and land plants (Wickett et al. 2014). Thus, species tree estimation methods that account for ILS, either explicitly or implicitly, are of interest.

An alternative to concatenation involves estimating gene trees (typically one per locus) and then applying a summary method. The most popular summary method to date, ASTRAL (Mirarab et al. 2014b), is a heuristic for the NP-hard Maximum Quartet Support Species Tree (MQSST) problem (Lafond and Scornavacca 2019), which can be framed as weighting quartets (four-leaf trees) by their frequencies in the input gene trees and then seeking a species tree *T* that maximizes the total weight of the quartets displayed by *T*. The optimal solution to MQSST is a statistically consistent estimator of the (unrooted) species tree under the MSC model, which is why heuristics for this problem are widely used in the context of multi-locus species tree estimation. Proofs of consistency typically assume the input gene trees are error-free (Roch et al. 2018); however, this is rarely the case. Gene trees estimated in recent studies have had low bootstrap support on average (Table 1 in Molloy and Warnow 2018), suggesting that gene tree estimation error (GTEE) is pervasive in modern phylogenomics data sets. GTEE has been shown to negatively impact the accuracy of summary methods in both simulation (e.g. Xi et al. 2015) and systematic studies (e.g. Meiklejohn et al. 2016). Together, GTEE and ILS present significant challenges to species tree estimation.

Scalability is also an issue when estimating species trees from large heterogeneous data sets. ASTRAL executes an exact (dynamic programming) algorithm for MQSST within a constrained version of the solution space constructed from the input gene trees. There have been many improvements to ASTRAL, with the latest version ASTRAL-III (Zhang et al. 2018) running in *O*((*nk*)^1.726^*x*) time, where *n* is the number of species (also called taxa), *k* is the number of gene trees, and *x* = *O*(*nk*) is the size of the constrained solution space. In addition, a recent method FASTRAL (Dibaeinia et al. 2021) runs ASTRAL-III in an aggressively constrained solution space to speedup species tree estimation. Importantly, the ASTRAL operates directly on the input set of *k* gene trees instead of explicitly constructing a set of Θ(*n*^4^) weighted quartets. This is in stark contrast to the other popular MQSST heuristics: weighted Quartet Max Cut (wQMC; Avni et al. 2014) and weighted Quartet Fiduccia-Mattheyses (wQFM; Mahbub et al. 2021).

The wQMC and wQFM methods take weighted quartets as input and thus require a preprocessing step, in which Θ(*n*^4^) quartets are weighted by the number of gene trees that display them. Both implement divide-and-conquer approach to species tree estimation, which is quite different than the approach used by ASTRAL. Interestingly, a recent study found wQFM outperforms ASTRAL-III under model conditions characterized by high ILS and high GTEE (Mahbub et al. 2021); however, the scalability of wQFM is limited due to the required preprocessing. In this paper, we enable improved accuracy and scalability under these challenging scenarios by introducing TREE-QMC.

## Results

### Overview of TREE-QMC Method

TREE-QMC builds upon the first widely-used MQSST heuristic, wQMC, which reconstructs the species tree in a divide-and-conquer fashion. At each step in the divide phase, an internal branch in the output species tree is identified; this branch splits the taxa into two disjoint subsets (Figure 1). The algorithm continues by recursion on the subproblems implied by the two subsets of taxa. Importantly, “artificial taxa” are introduced to represent the species on the opposite of the branch so that solutions to subproblems can be combined during the conquer phase. The recursion terminates when the subproblem has three or fewer taxa, as there is only one possible tree that can be returned (Supplementary Figure S1). At each step in the conquer phase, trees for complementary subproblems are connected at their artificial taxa, until there is a single tree on the original set of species.

**Figure 1:**
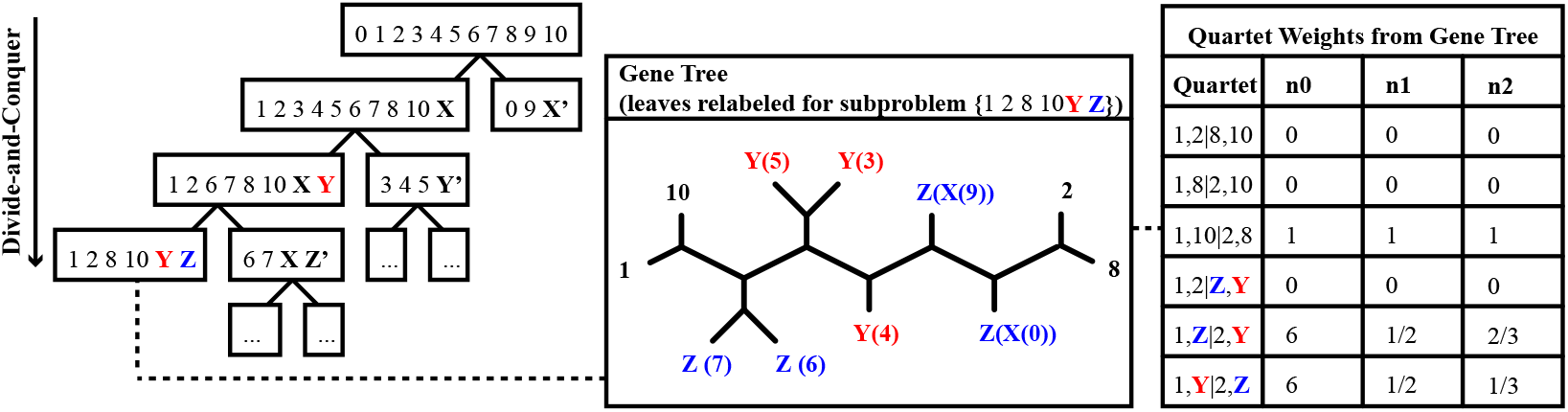
At each step in the divide phase, taxa are split into two disjoint subsets and then artificial taxa are introduced to represent the species on the other side of the split. To compute the quartet weights for a given subproblem, the leaves of each gene tree are relabeled by the artificial taxa. Without normalization (column n0), quartet 1, 2|*Y, Z* gets 0 votes and the alternative quartets get 6 votes each (note: quartet 1, *Y*|2, *Z* gets 6 votes by taking either species 5, 3, or 4 for label *Y* and either species 0 or 9 for label *Z*). With normalization, each gene tree gets one vote for each subset of four labels, although this vote can be split across the three possible quartets. In the uniform normalization scheme (column n1), we simply divide column n0 by the total number of votes cast in the unnormalized case. In the non-uniform normalization scheme (column n2), we leverage that structure implied by the divide phase of the algorithm; the idea is that species should have lesser importance each time they are re-labeled by artificial taxa.

Central to wQMC’s divide-and-conquer approach is a graph built from the (weighted) quartets. This graph is constructed in such a way that its max cut should correspond to a branch in the output species tree (Snir and Rao 2010, 2012; Avni et al. 2014). Our observation is that quartets with artificial taxa can have higher weights than quartets with only non-artificial taxa (called singletons) when looking at a single gene tree (Figure 1). As we will show, normalizing the quartet weights so that each gene tree gets one vote for every subset of four species greatly improves accuracy. The best performing normalization scheme (n2) weights quartets based on subproblem decomposition; specifically, quartets are upweighted if the species labeling their leaves are more closely related to the current subproblem (note: n1 denotes uniform normalization and n0 denotes no normalization). Moreover, we provide an algorithm to build the (normalized) quartet graph directly from the input gene trees, enabling TREE-QMC to have a time complexity of *O*(*n*^3^*k*) with some assumptions on subproblem sizes (see Methods section for details).

In the remainder of this section, we evaluate the performance of TREE-QMC (and its different normalization schemes) against the leading MQSST heuristics on simulated data. We then apply these methods to a real avian phylogenomics data set (Jarvis et al. 2014).

### Experimental Evaluation

We now give an overview of our simulation study; see Supplementary Materials for details.

#### Methods

TREE-QMC is compared against five leading MQSST heuristics: wQMC v1.3, wQFM v3.0, ASTRAL v5.5.7 (denoted ASTRAL-III or ASTRAL3), and FASTRAL. Two of these methods, wQMC and wQFM, which take weighted quartets instead of gene trees as input (the preprocessing step is performed using the script distributed on Github with wQFM). All methods are run in default mode. The current version of TREE-QMC requires binary gene trees as input so polytomies in the estimated gene trees are arbitrarily before running TREE-QMC (the same refinements are used in all runs of TREE-QMC to ensure a fair comparison across the normalization schemes).

#### Evaluation Metrics

All methods are compared in terms of species tree error, quartet score, and runtime. Species tree error is the percent Robinson-Foulds (RF) error (i.e., normalized RF distance between the true and estimated species trees multiple by 100). Because the true and estimated species tree are both binary, this quantity is equivalent to the percentage of false positive branches (i.e., internal branches in the estimated species tree that are incorrect and thus missing from the true species tree). Two-sided Wilcoxon signed-rank tests are used to evaluate differences between TREE-QMC-n2 versus FASTRAL as well as TREE-QMC-n2 versus ASTRAL3 (TREE-QMC-n2 is also compared against wQFM when possible). The quartet score is the number of quartets in the input gene trees that are displayed by the estimated species tree. All methods are run on the same data set on the same compute node, with a maximum wall clock time of 18 hours. The runtime of wQFM and wQMC includes the time to weight quartets based on the input gene trees (the fraction of time spent on this preprocessing phase is reported in the Supplementary Materials).

#### Simulated data sets

Our benchmarking study utilizes data simulated in prior studies, specifically the ASTRAL-II simulated data sets (Mirarab and Warnow 2015) as well as the avian and mammalian simulated data sets (Mirarab et al. 2014a). These data are generated by (1) taking a model species tree, (2) simulating gene trees within the species tree under the MSC, (3) simulating sequences down each gene tree under the GTR model, and (4) estimating a tree from set of gene sequences. Either the true gene trees from step 2 or the estimated gene trees from step 4 can be given as input to methods. This process is repeated for various parameter settings.

The avian and mammalian simulated data sets are generated from published species trees estimated for 48 birds (Jarvis et al. 2014) and 37 mammals (Song et al. 2012), respectively. The species tree branches are scaled to vary the amount of ILS, and the sequence length is changed to vary the amount of GTEE. There are 20 replicates for each model condition.

The ASTRAL-II data sets are generated from model species trees simulated under the Yule model given three parameters: species tree height, speciation rate, and number of taxa. The speciation rate is set so that speciation events are clustered near the root (deep) or near the tips (shallow) of the species tree. There are 50 replicates for each model condition (note that a new model species tree is simulated each replicate data set)

The data properties (ILS and GTEE levels) are summarized in Supplementary Tables S1 and S2. The ILS level is the percent RF error (between the true species tree and the true gene tree) averaged across all gene trees, and GTEE level is the percent RF error (between the true and estimated gene trees) averaged across all gene trees. Overall, these data sets cover a range of important model conditions. The results are presented in four experiments looking at the impact of varying the number of taxa, the species tree scale/height (proxy for ILS), the sequence length (proxy for GTEE), and the number of genes.

#### Number of Taxa

Figure 2 shows the impact of varying the number of taxa. The pipelines that need weighted quartets to be given as input (wQFM and wQMC) run on the order of seconds for 10 taxa, minutes for 50 taxa, and hours for 100 taxa, and did not complete within 18 hours (our maximum wallclock time) for the vast majority of data sets with 200 taxa. Importantly, the runtime of these pipelines is dominated by the time to weight Θ(*n*^4^) quartets by their frequency in the input gene trees (Supplementary Table S3). In contrast, TREE-QMC implements the same approach as wQMC but bypasses this preprocessing step, scaling to 1000 taxa and 1000 genes. For these data sets, FASTRAL, TREE-QMC-n2, and ASTRAL-III complete on average in 32 minutes, 64 minutes, and 5.3 hours, respectively (although ASTRAL-III fails to complete on 3/50 replicates within 18 hours). Thus, TREE-QMC-n2 is much faster than ASTRAL-III and is not much slower than FASTRAL. More importantly, TREE-QMC-n2 is significantly more accurate than either FASTRAL or ASTRAL-III when the number of taxa is 200 or greater. For these same conditions, quartet weight normalization, and especially the non-uniform (n2) scheme, improves TREE-QMC’s accuracy.

**Figure 2:**
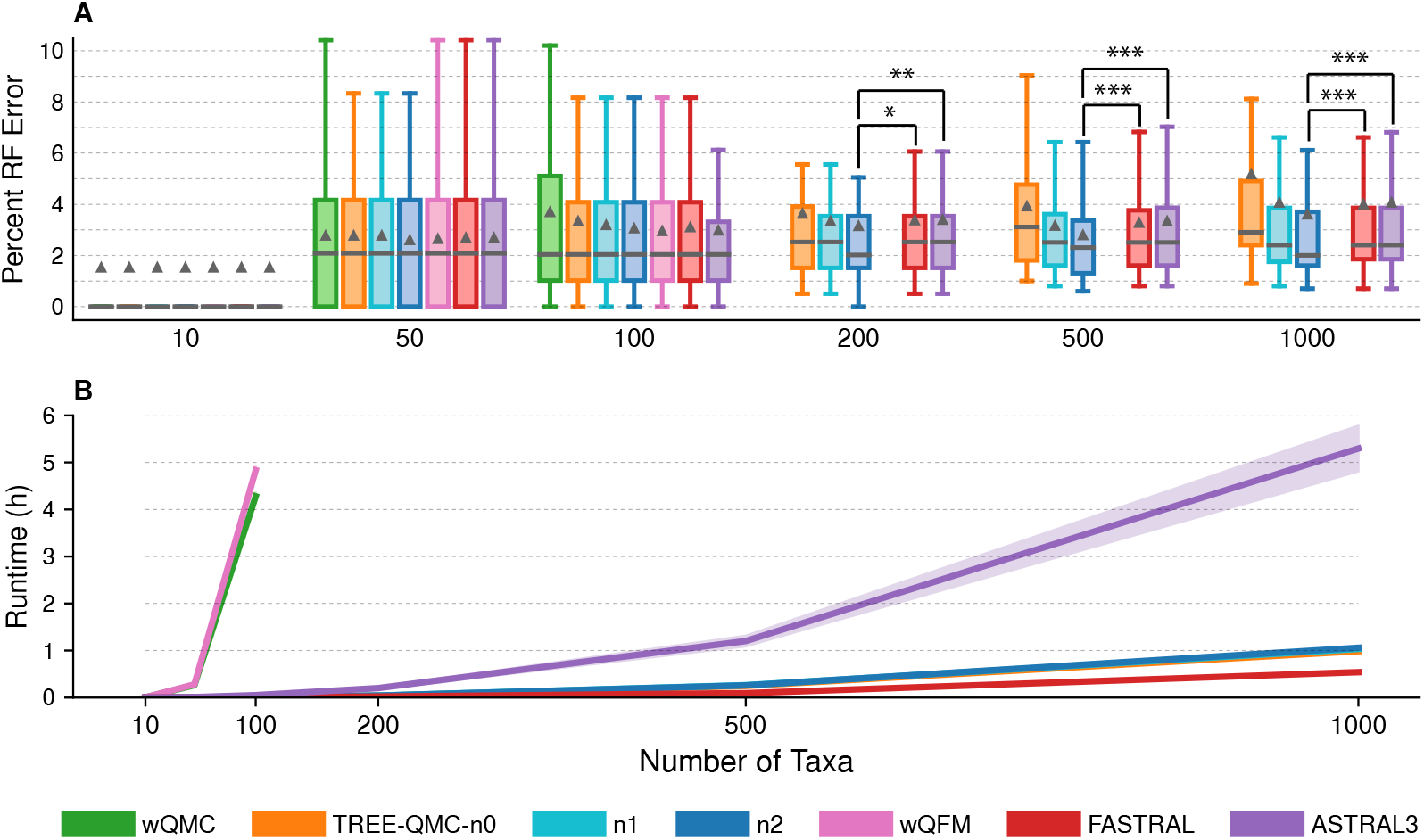
Impact of number of taxa. (A) Percent species tree error across replicates (bars represent medians; triangles represent means; outliers are not shown). The symbols *, **, and *** indicate significance at *p* < 0.05, 0.005, and 0.0005, respectively (all but * survive Bonferroni multiple comparison correction; see Supplementary Table S4 for details). (B) Mean runtime across replicates (shaded region indicates standard error). All data sets have species tree height 1X, shallow speciation, and 1000 estimated genes trees. The ILS level is 17–35% (ILS level), and GTEE level is 19–30%.

#### Incomplete Lineage Sorting (ILS)

##### ASTRAL-II data (200 taxa, 1000 estimated gene trees)

Figure 3 shows the impact of varying the species tree height and thus the amount of ILS for the ASTRAL-II data sets. TREE-QMC-n2, FASTRAL, and ASTRAL-III produce highly accurate species trees, with median species tree error at or below 6% for all model conditions (note that wQMC and wQFM cannot be run on these 200-taxon data sets within the maximum wall clock time). For some conditions, TREE-QMC-n2 is significantly more accurate than FASTRAL or ASTRAL-III, and there is no significant difference between pairs of methods for the other conditions. Notably, quartet weight normalization improves the accuracy of TREE-QMC; this effect is most pronounced when the amount of ILS was very high (species tree height: 0.5X). On these same conditions, ASTRAL-III is much slower than the other methods, taking taking 73 minutes on average for the highest amount of ILS (species tree height: 0.5X) compared to 5 minutes on average for the lowest amount of ILS (species tree height: 5X). In contrast, both TREE-QMC-n2 and FASTRAL are quite fast, taking on average less than 3 minutes for model conditions with 200 or fewer taxa.

**Figure 3:**
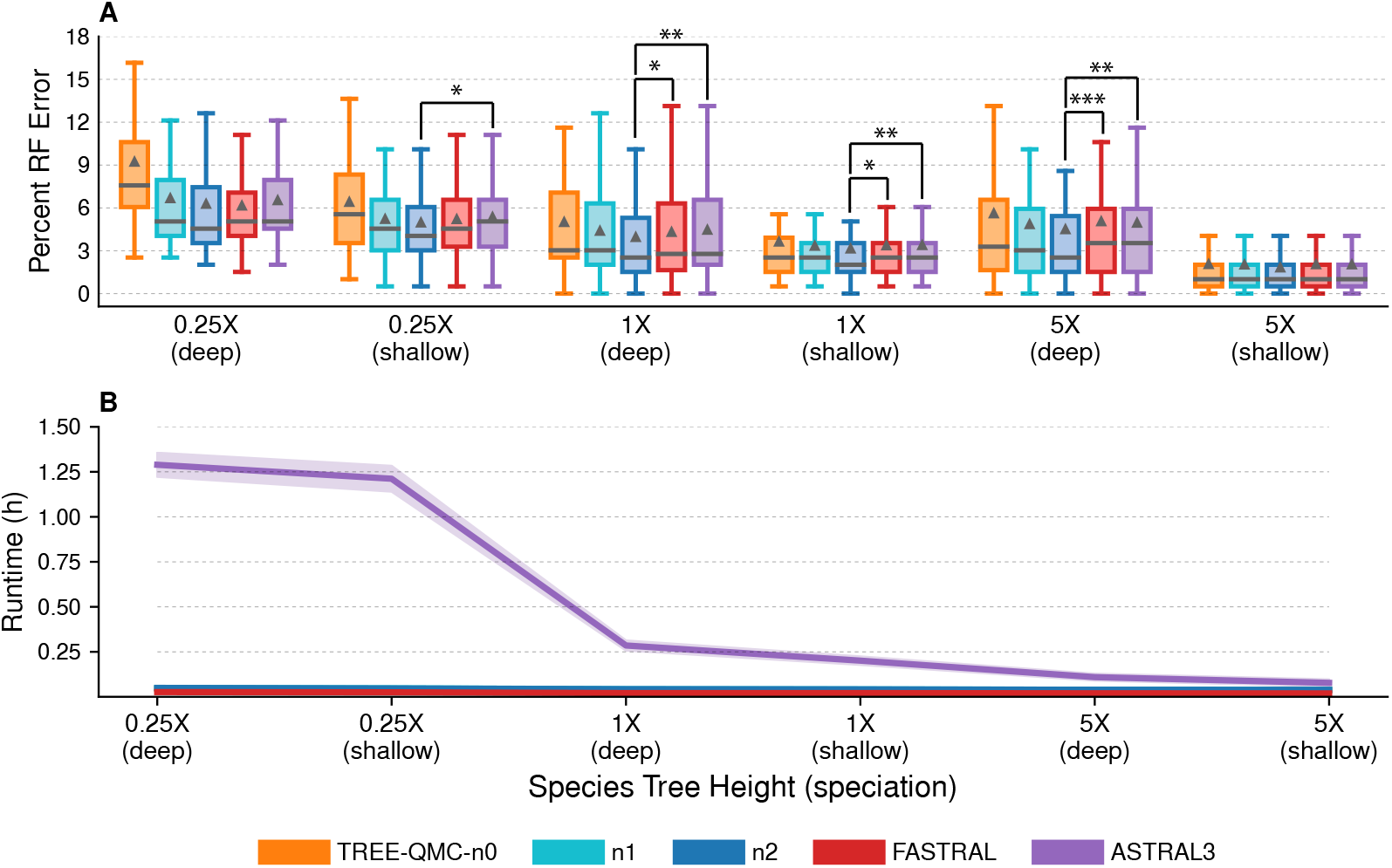
Impact of the amount of ILS on MQSST heuristics. (A) Percent species tree error across replicates (bars represent medians; triangles represent means; outliers are not shown). The symbols *, **, and *** indicate significance at *p* < 0.05, 0.005, and 0.0005, respectively (three tests survive multiple comparison corrections; see Supplementary Table S5 for details). (B) Mean runtime across replicates (shaded region indicates standard error). All data sets have 200 taxa and 1000 estimated gene trees. One model condition with species tree height 1X and shallow speciation is repeated from Figure 2. For species tree heights 0.5X, 1X, and 5X, the ILS level is 68–69%, 34%, and 9–21%, respectively, and the GTEE level is 44%, 27%–34%, and 21-28%, respectively.

##### Avian simulated data (48 taxa, 1000 estimated gene trees)

Figure 4A–C shows the impact of varying the species tree scale and thus ILS on the avian simulated data sets. The original wQMC method is the least accurate method and is even less accurate than TREE-QMC-n0 (no normalization). Normalization improves the performance of TREE-QMC for these data, enabling TREE-QMC-n2 to be among the most accurate methods when the amount of ILS is higher (species tree scales: 0.5X and 1X). Testing for differences between TREE-QMC-n2 versus wQFM, FASTRAL, and ASTRAL-III reveals that either TREE-QMC-n2 is significantly better or there are no significant differences between the pairs of methods. All methods finish quickly: wQMC and wQFM completes in less than 13 minutes on average, ASTRAL-III completes in less than 4 minutes on average, and the other methods finish in less than 1 minute on average.

**Figure 4:**
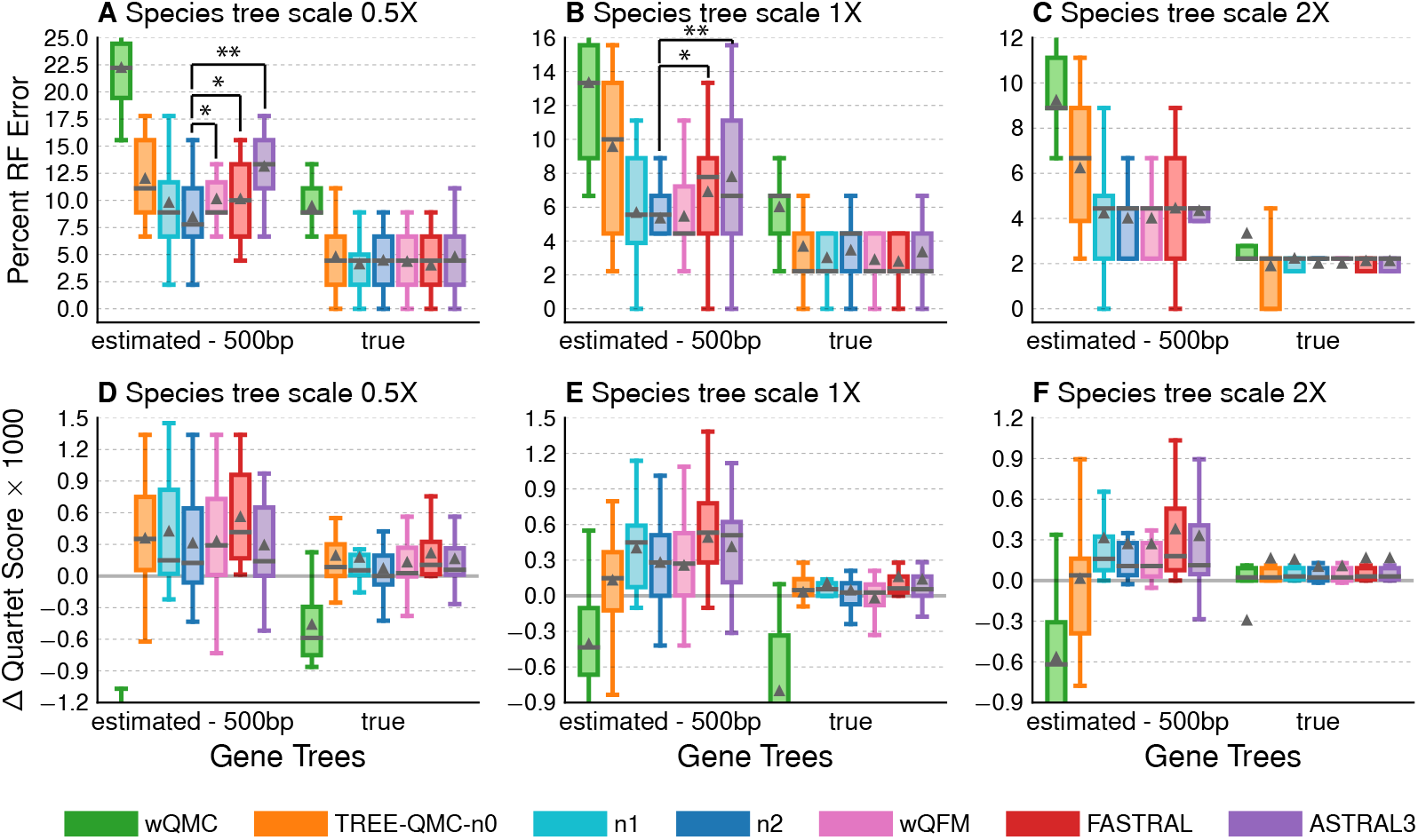
Impact of ILS and GTEE on MQSST heuristics. (A), (B, and (C) Percent species tree error for the avian data set with 1000 estimated or true gene trees and species tree scales 0.5X, 1X, and 2X, respectively. Two-sided Wilcoxon-signed ranked tests were used to evaluate differences between TREE-QMC-n2 versus wQFM, FASTRAL, and ASTRAL3 (9 tests per subfigure). The symbols *, **, and *** indicate significance at *p* < 0.05, 0.005, and 0.0005, respectively (for 0.5X species tree scale with estimated gene trees, the difference between TREE-QMC-n2 and ASTRAL-II survives Bonferroni multiple comparison correction; see Supplementary Table S6 for details). (D), (E), and (F) show the quartet score for the estimated species tree minus the quartet score for the true species tree times 1000 for species tree scales 0.5X, 1X, and 2X, respectively. For species tree heights 0.5X, 1X, and 2X, the ILS level is 60%, 47%, and 35%, respectively, and the GTEE level is 60%, 60%, and 62%, respectively. Results for wQMC are cut off because otherwise the trends cannot be observed (see Supplementary Figure S2 for full y-axes).

Figure 4D–F shows the difference between the quartet score of the estimated species tree minus the quartet score of the true species tree (species trees were scored with the same gene trees used to estimate them). Higher quartet scores do not necessarily correspond to greater accuracy. For example, TREE-QMC-n0 is always less accurate than TREE-QMC-n2 but the former a higher quartet score for the lowest ILS level (Figure 4D) and a lower quartet score for the middle ILS level (Figure 4E). In general, the best performing methods find species trees with higher quartet scores than the true species tree when gene trees have high estimation error.

##### Mammalian simulated data (37 taxa, 200 estimated gene trees)

All methods have similar performance for the mammalian data, although these data sets represent easier model conditions in terms of ILS and GTEE levels (Supplementary Figure S3, Supplementary Table S5).

#### Gene Tree Estimation Error (GTEE)

##### Avian simulated data (48 taxa, 1000 gene trees)

Figure 4A–C also shows the impact of GTEE for each species tree scale (ILS level). Across all ILS levels, methods are either given true gene trees or estimated gene trees with substantial error (60-62%). Without GTEE, there is no significant differences between TREE-QMC-n2 versus the other leading methods (wQFM, FASTRAL, and ASTRAL-III), and all versions of TREE-QMC perform similarly so the utility of normalization is diminished. In addition, methods find species trees with similar quartet scores to the true species tree when given true gene trees as input. Lastly, the performance of wQMC is inline with the other methods (Figure 4C) when there is very little gene tree heterogeneity due to ILS or GTEE.

##### Mammalian simulated data (37 taxa, 200 gene trees)

Similar trends between methods are observed for mammalian simulated data sets when varying the sequence lengths (Supplementary Figure S4). TREE-QMC is significantly more accurate than FASTRAL and ASTRAL-III for the shortest sequence length (250 bp; GTEE level 43%) and there are no differences between methods otherwise.

#### Number of Genes

Similar trends between methods are observed for varying the number of genes (Supplementary Figures S5). Overall, TREE-QMC-n2 is the best performing method, with error rates similar to wQFM (although, as shown in the first experiment, TREE-QMC-n2 scales to data sets with larger numbers of taxa).

### Avian phylogenomics data set

We also re-analyze the avian data set from Jarvis et al. (2014) with 3,679 ultraconserved elements (UCEs). This data set includes the best maximum likelihood tree and the set of 100 bootstrapped trees for each UCE. Although the true species tree is unknown, we discuss the presence and absence of strongly corroborated clades, such as Passerea and six of the magnificent seven clades excluding clade IV (Braun and Kimball 2021). We also compare methods to the published concatenation tree estimated by running RAxML (Stamatakis 2014) on UCEs only (Jarvis et al. 2014); thus the comparison between concatenation and the MQSST heuristics is on the same data set. Branch support is computed for the estimated species trees using ASTRAL-III’s local posterior probability (Sayyari and Mirarab 2016) as well as using multi-locus bootstrapping (MLBS) (Seo 2008). We repeat this analysis (except MLBS) on the TENT data (14,446 gene trees), which includes gene trees estimated on UCEs as well as exons and introns. In this case, methods are compared to the published TENT concatenation tree estimated by running ExaML (Kozlov et al. 2015).

### UCE data

For the UCE data (48 taxa, 3679 gene trees), ASTRAL-III complete in 65 minutes, making it the most time consuming method. All other methods run in less than a minute; however, the preprocessing step to weight quartets for wQFM takes 41 minutes.

FASTRAL and ASTRAL-III produce the same species tree, and TREE-QMC-n2 and wQFM produce the species tree. We compare these two trees to the published concatenation tree for UCEs (Figure 5). There are many similarities between these three trees, as all contain the magnificent seven clades. The TREE-QMC-n2 and FASTRAL trees differ from the concatenation tree by 7 and 9 branches, respectively, putting the TREE-QMC-n2 tree slightly closer to the concatenation tree than the FASTRAL tree. Notably, the TREE-QMC-n2 tree recovers Passerea and Afroaves and fails to recover Columbea, like the concatenation tree and unlike the ASTRAL-III tree (note that Passerea was considered to be strongly corroborated, after accounting for data type effects, by Braun and Kimball 2021). Overall, there are only five branches that differ between the TREE-QMC-n2 tree and the FASTRAL tree; all of these branches have nearly equal quartet support for their alternative resolutions so that both trees represent reasonable hypotheses.

**Figure 5:**
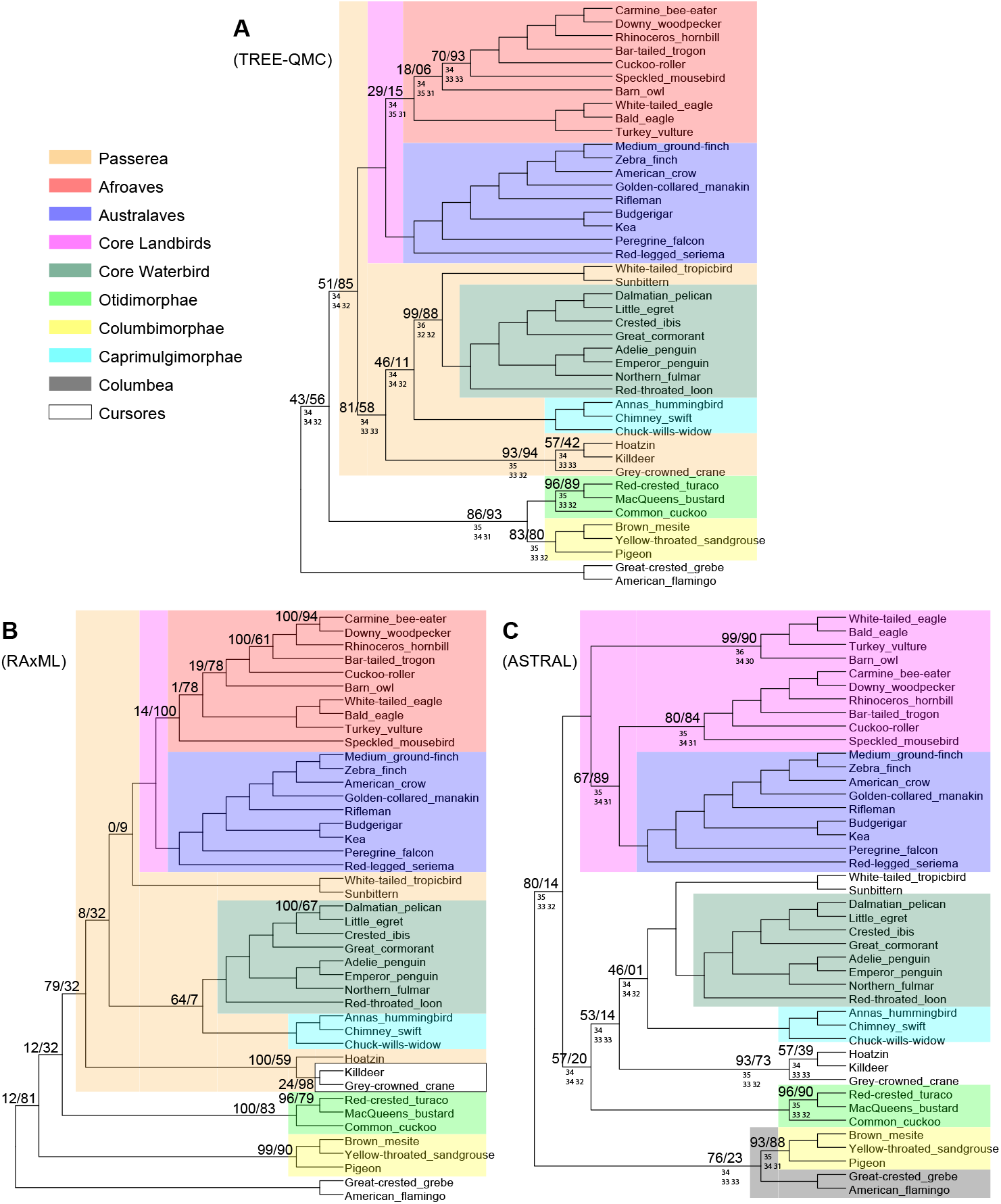
(A) Species tree estimated from UCE gene trees using TREE-QMC-n2 or wQFM. (B) Species tree estimated from concatenated UCE alignment using RAxML. (C) Species tree estimated from UCE gene tres using ASTRAL-III or FASTRAL. Above the branch, we show support values *X/Y*, where *X* is estimated using ASTRAL’s local posterior probability (multiplied by 100) and *Y* is using MLBS for subfigures A and C and *Y* is the bootstrap support computed by RAxML for subfigure B. Support values are only shown when *X* is less than 100. Below the branch, we show the quartet support (the two values below it correspond to quartet support for the two alternative resolutions of the branch). Taxa outside of Neoaves are not shown as all methods recovered the same topology outside of Neoaves.

### TENT data

For the TENT data (48 taxa, 14446 gene trees), TREE-QMC-n2 and FASTRAL complete in less than 3 minutes, whereas it takes 2.35 hours to weight quartets. wQFM completes in less than a minute after this preprocessing phase. We do not run ASTRAL-III as this analysis was reported to take over 30 hours (Dibaeinia et al. 2021).

All three methods produce a different tree, which is compared to the published concatenation tree for TENT data (Supplementary Figure S6). None of the trees recover Passera, and only the concatenation and wQFM trees recover Afroaves, although this branch has very local support (local posterior probability of 0.0) in the wQFM tree. Once again, the TREE-QMC-n2 and wQFM trees are closest to the concatenation tree, with the TREE-QMC-n2, wQFM, and FASTRAL trees differing from it by 8, 8, and 10 branches, respectively. There are 5 branches that differ between the wQFM tree and the TREE-QMC-n2 tree (notably two of these branches in the wQFM have very low support: local posterior probability of 0.03 and 0.0). There are only 3 branches that differ between the TREE-QMC-n2 tree and the FASTRAL tree; as with the UCE data, these branches are reasonable based on quartet support for their alternative resolutions.

## Discussion

Our method TREE-QMC builds upon the algorithmic framework of wQMC (Avni et al. 2014) by introducing the *normalized* quartet graph and showing that it can be computed directly from gene trees. These contributions together enable our new method TREE-QMC to be highly competitive with the leading MQSST heuristics, even outperforming them. In our simulation study, TREE-QMC (with non-uniform normalization) is more accurate than other methods when the amount of gene tree heterogeneity due to ILS and/or GTEE is high and when the number of species is large. These scenarios are known challenges to species tree estimation and the issue of GTEE, in particular, has motivated a new version of ASTRAL, dubbed weighted ASTRAL (Zhang and Mirarab 2022), which was published during our study. The idea behind weighted ASTRAL is that quartets should be weighted based on the estimated gene trees, specifically branch support on the internal branch of the quartet and/or branch lengths on the terminal edges of the quartet. TREE-QMC’s non-uniform normalization scheme also weights quartets but does so based subproblem division (i.e., quartets are upweighted if they are on species in more closely related subproblems, which ideally reflects closeness in the true species tree). In the future, it would be interesting to compare TREE-QMC to weighted ASTRAL as well as to implement other quartet weighting schemes within TREE-QMC.

There are several other opportunities for future work worth mentioning. First, the version of TREE-QMC presented here requires binary gene trees as input. Thus, TREE-QMC was given gene trees that are randomly refined in our experimental study, whereas all other methods were given gene trees with polytomies. This did not have a negative impact on TREE-QMC’s performance relative to the other MQSST heuristics; however, it would be worth exploring this issue further. Ultimately, this inherent limitation of TREE-QMC could be addressed by devising an efficient algorithm for computing the “ edges” in the quartet graph (see Methods section), although this would come at the cost of increased runtime. Second, the experimental study presented here only evaluates TREE-QMC in the context of multi-locus species tree estimation where gene tree can be discordant with the species tree due to ILS and/or GTEE. Our study does not address the use of TREE-QMC as a more general quartet-based supertree method, and future work should explore whether quartet weight normalization is beneficial in this context. Lastly, TREE-QMC’s algorithm operates on gene trees that are multi-labeled due to artificial taxa, so the algorithms presented here can be applied to gene trees that are multi-labeled due to other causes, such as multiple individuals being sampled per species (Rabiee et al. 2019) or genes evolving via duplications (Legried et al. 2021; Zhang et al. 2020; Yan et al. 2021; Smith et al. 2022). Future work should explore the effectiveness of TREE-QMC under these conditions as well those characterized by missing data due to gene loss or other causes (Nute et al. 2018).

## Methods

We begin with some notation and terminology for phylogenetic trees. A *phylogenetic tree T* is a triplet (*g*, 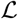, *ϕ*), where *g* is a connected acyclic graph, 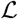 is a set of labels (species), and *ϕ* maps leaves in *g* to labels in 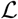. If *ϕ* is a bijection, we say that *T* is *singly-labeled*; otherwise, we say *T* is *multi-labeled*. Trees may be either *unrooted* or *rooted*. Henceforth, all trees are *binary*, meaning that non-leaf, non-root vertices (referred to as *internal* vertices) have degree 3. For a tree *T*, we denote its edge set as *E*(*T*), its internal vertex set as *V*(*T*), and its leaf set as *L*(*T*). Edges in an unrooted tree are undirected, whereas edges in a rooted tree are directed away from the root, a special vertex with in-degree 0 (all other vertices have in-degree 1). To transform an unrooted tree *T* into a rooted tree *T_r_*, we select an edge in *T*, sub-divide it with a new vertex *r* (the root), and then orient the edges of *T* away from the root. Conversely, we transform a rooted tree *T_r_* into an unrooted tree *T* by undirecting its edges and then suppressing any vertex with degree 2. Sometimes we consider a phylogenetic tree *T restricted* to a subset of its leaves *R* ⊆ *L*(*X*). Such a tree, denoted *T*|*_R_*, is created by deleting leaves in *L*(*T*) \ *R* and suppressing any vertex with degree 2 (while updating branch lengths in the natural way).

To present TREE-QMC, we need two additional concepts: *bipartitions* and *quartets*. A bipartition splits a set 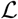 of labels into two disjoint sets: 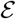 and 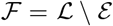. Each edge in a (singly-labeled, unrooted) tree *T* induces a bipartition because deleting an edge creates two rooted subtrees whose leaf labels form the bipartition 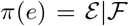. A given bipartition is displayed by *T* if it is in the set {*π*(*e*): *e* ∈ *E*(*T*)}. The bipartition is trivial if 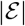 or 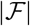 is 1; otherwise, it is non-trivial. A quartet *q* is an unrooted, binary tree with four leaves *a, b, c, d* labeled by *A, B,C, D*, respectively. It is easy to see that there are three possible quartet trees given by their one non-trivial bipartition: *a, b*|*c, d, a, c*|*b, d*, and *a, d*|*b, c* (note that we typically use lower case letters to denote leaf vertices and capital letters to denote leaf labels, although this distinction is only important when trees are multi-labeled). A set of quartets can be defined by a unrooted tree *T* by restricting *T* to every possible subset of four leaves in *L*(*T*); the resulting set *Q*(*T*) is referred to as the quartet encoding of *T*. If *T* is multi-labeled, then some of the quartets in *Q*(*T*) will have multiple leaves labeled by the same label. Lastly, we say that *T* displays a quartet *q* if *q* ∈ *Q*(*T*).

### Review of wQMC

As previously mentioned, our new MQSST heuristic, TREE-QMC, builds upon the divide-and-conquer method wQMC (Avni et al. 2014). To produce a bipartition on 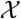, wQMC constructs a graph from 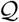, referred to as the **quartet graph**, and then seeks its maximum cut (Snir and Rao 2010, 2012; Avni et al. 2014). The quartet graph is formed from two complete graphs, 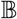 and 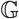, both on vertex set *V* (i.e., there exists a bijection between *V* and 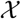). All edges in 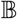 and 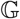 are initialized to weight zero. Then, each quartet 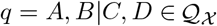 contributes its weight 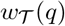 to two “bad” edges in 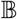 and four “good” edges in 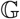, where 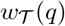 corresponds to the number of gene trees in the input set 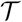 that display *q*. The bad edges are based on sibling pairs: (*A, B*) and (*C, D*). The good edges are based on non-sibling pairs: (*A, C*), (*A, D*), (*B, C*), and (*B, D*). We do not want to cut bad edges because siblings should be on the same side of the bipartition; conversely, we want to cut good edges because non-siblings should be on different sides of the bipartition. Ultimately, we seek a cut 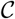 to maximize 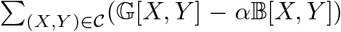, where *α* > 0 is a hyperparameter that can be optimized using binary search. Although MaxCut is NP-complete (Karp 1972), fast and accurate heuristics have been developed (Dunning et al. 2018). The cut gives a bipartition in the output species tree and the wQMC method proceeds by recursion on the two subsets of species on each side of the bipartition. Artificial taxa are introduced to represent the species on the other side of the bipartition.

### Quartet Weight Normalization

Our key observation is that artifical taxa change the quartet weights so that a single gene tree will vote multiple times for quartets on artificial taxa and only once for quartets on only non-artificial taxa (called singletons). As shown in Figure 1, the weight of quartet *M, N*|*O, P* is

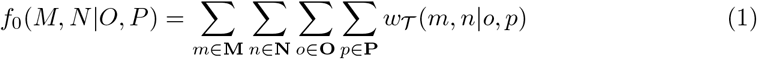

where 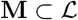 denotes the set of leaves (i.e., species) in *T* associated with label *M* (and and similarly for **N**, **O**, **P**). When labels *M, N, O, P* are all singletons, each gene tree casts exactly one vote for one of the three possible quartets: *M, N*|*O, P* or *M,O*|*N,P* or *M,P*|*N, O* (assuming no missing data). Otherwise, each gene tree casts |**M**| · |**N**| · |**O**| · |**P**| votes (again assuming no missing data) and thus can vote for more than one topology.

We propose to normalize the quartet weights so that each gene tree casts one vote for each subset of four labels, although it may split its vote across the possible quartet topologies in the case of artificial taxa. In the simplest case, we simply divide by the number of votes cast so the weight of *M, N*|*O, P* becomes

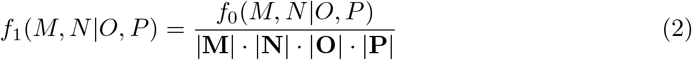

This can be implemented efficiently by assigning an importance value *I*(*x*) to each species 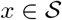 and then compute the weight as

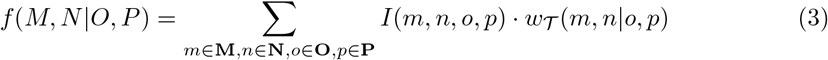

where *I*(*m, n, o, p*) = *I*(*m*) · *I*(*n*) · *I*(*o*) · *I*(*p*). Specifically, Equation 3 reduces to Equation 2 when *I*(*m*) = |**M**|^-1^ for all *m* ∈ **M** (and similarly for **N**, **O**, **P**). Because all species with the same label are assigned the same importance value, we refer to this approach as *uniform normalization (n1)*. More broadly, the quartet weights will be normalized whenever Equation 3 corresponds to a weighted average, meaning that

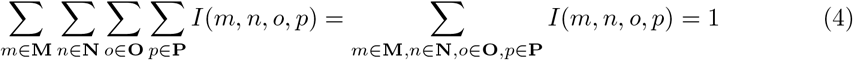

It is easy to see that this will be the case whenever ∑_*m*∈*M*_ *I*(*m*) = 1 (and similarly for **N**, **O**, **P**). Note that in *unnormalized (n0)* case, we assign all species an importance value of 1 so that Equation 3 reduces to Equation 1.

We now describe how to normalize quartet weights while leveraging the hierarchical structure implied by artificial taxa by assigning importance values to species with the same label. The idea is that species should have lesser importance each time they are *re-labeled* by an artificial taxon. In Figure 1, artificial taxon *Z* represents species **Z** = {0, 6, 7, 9} but species 0 and 9 were previously labeled by artificial taxon *X*. This relationship can be represented as the rooted “phylogenetic” tree *T_Z_* given by newick string: (6, 7, (0, 9)*X*)*Z*. We use *T_Z_* to assign importance values to all species *z* ∈ **Z**, specifically

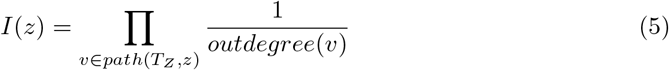

where *outdegree*(*v*) is the out-degree of vertex *v* and *path*(*T_Z_, z*) contains the vertices on the path in *T_Z_* from the root to the leaf labeled *z*, excluding the leaf. Continuing the example, 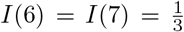 and 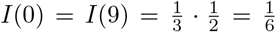. By construction, ∑_*z*∈**Z**_ *I*(*z*) = 1 so this approach normalizes the quartet weights. Because different species with the same label can have different weights, we refer to this approach *non-uniform normalization (n2)*. In our experimental study, normalizing the quartet weights in this fashion improved species tree accuracy for challenging model conditions.

### Efficient Quartet Graph Construction

We now describe our approach for constructing the quartet graph directly from the input gene trees, which is implemented within our new method TREE-QMC. The total weight of bad edges between *X* and *Y*, denoted 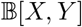, is the number of quartets (displayed by the input gene trees) with *X*, *Y* as siblings (and similarly for 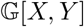 but non-siblings). Note that these quantities can be computed by summing over the number of bad and good edges contributed by each gene tree *T*. Henceforth, we consider how to compute 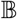 and 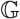 for a single gene tree.

We begin by considering a singly-labeled, binary gene tree *T* with *n* leaves. In this case, we can compute the number of good edges between *X, Y* via 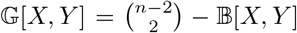, where n is the number of leaves in *T*. Because *T* is singly-labeled, there is exactly one leaf associated with label *X*, denoted *x*, and one leaf associated with label *Y*, denoted *y*. To compute 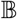 efficiently, we consider the unique path connecting leaves *x* and *y* in *T* (Figure 6a). Deleting the edges on this path (and their end points) produces a forest of *K* rooted subtrees, denoted {*t*_1_, *t*_2_,…, *t_K_*}. Let *w* and *z* be two leaves of subtrees *t_i_* and *t_j_*, respectively. Then, *T* displays quartet *x, w*|*z, y* for *i* < *j*, quartet *x, y*|*w, z* for *i* = *j*, and quartet *x, z*|*w, y* for *i* > *j*. To summarize, *x, y* are siblings if and only if leaves *w, z* are in the same subtree off the path from *x* to *y*. It follows that 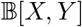 can be computed by considering all ways of selecting two other leaves from the same subtree for all subtrees on the path from *x* to *y*.

**Figure 6:**
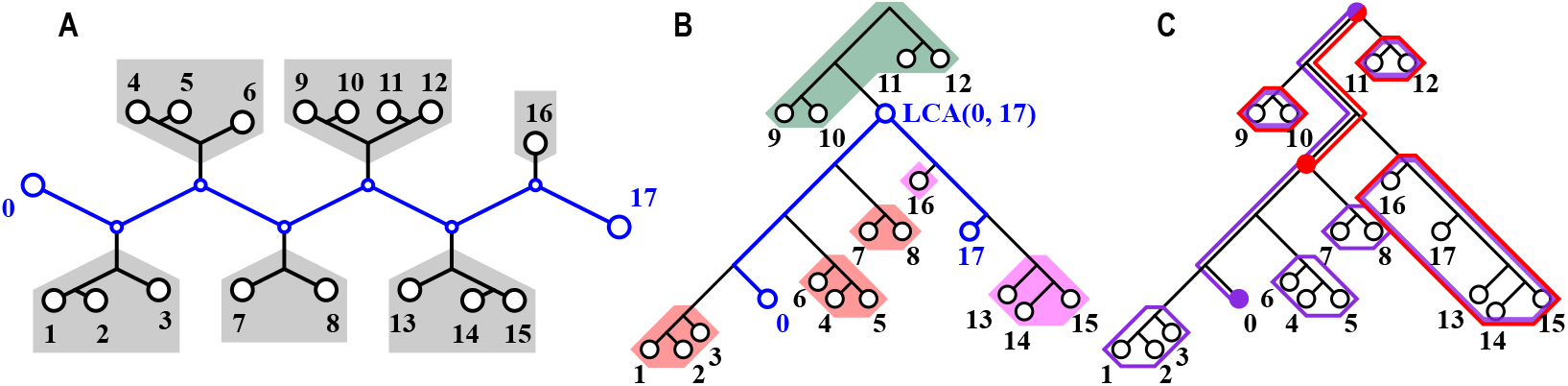
To count the quartets induced by *T* with 0 and 17 as siblings, we consider the path between them (shown in blue in (a)). The deletion of the path produces 6 rooted subtrees (highlighted in grey). Because 0 and 17 are siblings in a quartet if and only if the other two taxa are drawn from the same subtree, the number of bad edges can be computed as 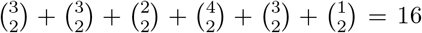. Here we show how to compute the number of quartets induced by *T* with 0 and 17 as siblings after rooting *T* arbitrarily. Subfigure (b) shows that we need to consider the number of ways of selecting two taxa from the same subtree for three cases: (1) the subtree above the *lca*(0, 17) (highlighted in green), (2) all subtrees off the path from the *lca*(0, 17) to the left taxon 0 (highlighted in red), and (3) all subtrees off the path from the *lca*(0, 17) to the right taxon 17 (highlighted in pink). Case 1 can be computed in constant time if we know the number of leaves below the LCA, that is, 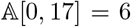 (Eq. 8). Cases 2 and 3 can also be computed in constant time as follows. Subfigure (c) shows the prefix of the left child of the *lca*(0, 17), denoted *p*[*lca*(0, 17).*left*] is the number of ways of selecting two taxa from the same subtree for all subtrees circled in red, which are off the path from the root to this vertex. Similarly, the the prefix of taxon 0, denoted *p*[0], is the number of ways of selecting two taxa from the same subtree for all subtrees circled in blue, which are off the path from the root to 0. Therefore, the number of ways of selecting two taxa from all subtrees in case 2 (i.e., subtrees highlighted in red in subfigure (b)) is 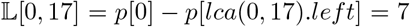 (Eq. 9). Case 3 (not shown) can be computed as 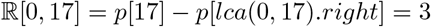 (Eq. 10). Putting this all together gives 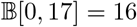 (Eq. 6).

This observation can be used to count the quartets efficiently when gene trees are singly-labeled. However, we need to be more careful when *T* is multi-labeled, which is typically the case due to artificial taxa. Following our example, suppose that we want to count the number of bad edges between 0 and 17 contributed by the subtree with leaves 4, 5, and 6. However, if leaves 4 and 5 are both re-labeled by artificial taxon *M*, the quartet on 0, 17|4, 5 corresponds to quartet 0, 17|*M, M* has no topological information and should not be counted. The other quartets 0, 17|4, 6 and 0, 17|5, 6 correspond to 0, 17|*M*, 6 and thus should be counted.

We now present an algorithm for computing 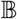 in *O*(*s*^2^*n*) time, where n is the number of leaves in gene tree *T*, and *s* is the number of labels in the subproblem (henceforth we let a denote the number of singletons and *b* denote the number of artificial taxa so the subproblem size is *s* = *a* + *b*). Our approach breaks down the calculation into three cases:

1. *X, Y* are both singletons,
2. *X* is a singleton and *Y* is an artificial taxon (or vice versa), and
3. *X, Y* are both artificial taxa.

To summarize our results, *B*[*X, Y*] can be computed for all pairs *X, Y* in case 1, case 2, and case 3 in *O*(*a*^2^) time, *O*(*abn*) and *O*(*b*^2^*n*) time, respectively. Thus, we can construct the quartet graph from k gene trees in *O*(*s*^2^*nk*) time (Theorem 1 in the Supplementary Materials). Afterwards, we seek a max cut using an *O*(*s*^3^) heuristic implemented in the open source library MQLib (Dunning et al. 2018). This gives us the final runtime of *O*(*s*^2^*nk* + *s*^3^) for each subproblem. If the division into subproblems is perfectly balanced, the divide- and-conquer algorithm runs in *O*(*n*^3^*k*) time (Theorem 2 in the Supplementary Materials). Although we do not expect perfectly balanced subproblems in practice, we found TREE-QMC to be fast in our experiments.

### Computing the number of bad edges given a singly-labeled gene tree

We first present an algorithm for computing the number of bad edges given a singly-labeled gene tree *T*. After rooting *T* arbitrarily, we again consider the path between *x* and *y*, which now goes through their lowest common ancestor, denoted *lca(x, y*) (Figure 6b). This allows us to break the computation into three parts

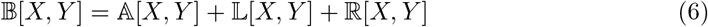

where 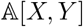 is the number of ways of selecting two leaves from the subtree above *lca*(*x, y*), 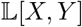 the number of ways of selecting two leaves from the same subtree for all subtrees off the path from *lca*(*x, y*) to leaf in it’s *left* subtree (say x), and 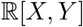 the number of ways of selecting two leaves from the same subtree for all subtrees off the path from *lca*(*x, y*) to the leaf in its *right* (say *y*). As we will show, each of these quantities can be computed in constant time, after an *O*(*n*) preprocessing phase, in which we compute two values for each vertex *v* in *T*. The first value *c*[*v*] is the number taxa below vertex *v*. The second value *p*[*v*], which we refer to as the “prefix” of *v*, is the number of ways to select two taxa from the same subtree for all subtrees off the path from the root to vertex *v* (Figure 6c). It is easy to see that *c* can be computed in *O*(*n*) time via a post-order traversal. After which, *p* can be computed in *O*(*n*) via a preorder traversal, setting

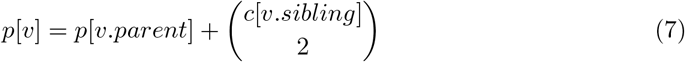

after initializing *p*[*root*] = 0. Now we can compute the quantities:

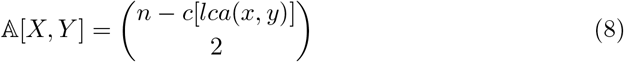

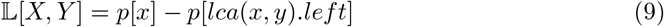

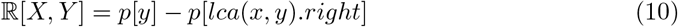

where *v.left* denotes the left child of *v* and *v.right* denotes the right child of *v* (see Figure 6c). It is possible to access *lca*(*x, y*) in constant time after *O*(*n*) preprocessing step (Gusfield 1997), although we implemented this implicitly by computing the entries of 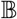 during a post-order traversal of *T*. Thus, we can compute 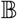 in *O*(*n*^2^) time, provided that T is singly-labeled.

### Computing the number of bad edges given a multi-labeled gene tree

We now present an algorithm for computing the number of bad edges 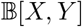 given a multi-labeled gene tree *T*. As previously mentioned, this breaks down into three cases. The first case (*X, Y* are both singletons) is below and the remaining two cases are presented in the Supplementary Materials.

Again, we focus on the number of ways to select two leaves *w, z* from a collection of subtrees. When *T* is multi-labeled, it is possible for two leaves *w, z* to have the same label. Thus, we now need to count the number of ways to select two leaves *z, w* below vertex u so that they are **uniquely labeled** *Z* ≠ *W* (note that we use capital letters *W* and *Z* to denote the current labels of leaves *w* and *z*, respectively). This modified binomial is computed by revising the preprocessing phase. We now let *c*_0_[*v*] denote the number of leaves labeled by singletons below vertex *v* and let *c_D_*[*v*] denote the number of leaves labeled by artificial taxon *D* below vertex *v*. Thus, for each vertex *v*, we store a vector *c*[*v*] of length *b* + 1, where *b* is the number of artificial taxa in *T*. As before, we can compute *c* in *O*(*bn*) time via a postorder traversal. However, the number of ways to select two leaves with different labels is now broken into three cases:

1. the number of ways to select two singletons, which equals 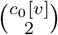,
2. the number of ways to select one singleton and one artificial taxa, which equals 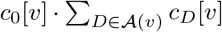, where 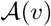 is the set of artificial taxa below vertex *v*, and
3. the number of ways to select two artificial taxa, which equals 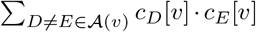.

Putting this all together gives the **modified binomial coefficient**:

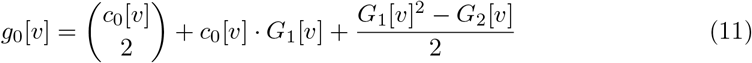

where 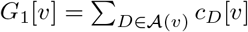 and 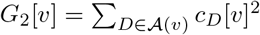. At each vertex, the calculation of *G*_1_[*v*] and *G*_2_[*v*] takes *O*(*b*) time, after which we can compute *g*_0_[*v*] in constant time. Thus, *g*_0_ can be computed in *O*(*bn*) time. Note that we also need to compute modified binomial coefficient for the subtree “above” vertex *v*, denoted *g*_0_[*v.above*]. This can be computed in a similar fashion by noting that number of singletons above *v* is *a* – *c*_0_[*v*] and that the number of leaves above *v* labeled by each artificial taxon *D* is |**D**| – *c_D_*[*v*].

Using the modified binomial, we can apply our algorithm for singly-labeled trees by redefining prefix sum:

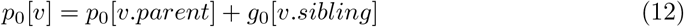

and then redefining the quantities from which we can compute *B*[*x, y*] in constant time, that is, 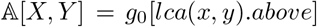, and 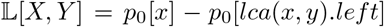, and 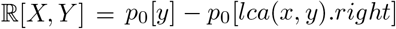. As there are *a*^2^ pairs of singletons in the subproblem, the total runtime is *O*(*a*^2^ + *bn*).

### Normalizing quartet weights when computing bad edges

To normalize the quartet weights, 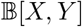 becomes the *weighted* sum of quartets with *X, Y* are siblings, where each quartet *x, y*|*z, w* is weighted by *I*(*x, y, z, w*) = *I*(*x*)*I*(*y*)*I*(*z*)*I*(*w*), where *I*(*x*) is the importance value assigned to leaf *x* (which corresponds to a species in the singly-labeled gene tree). When *X, Y* are singletons,

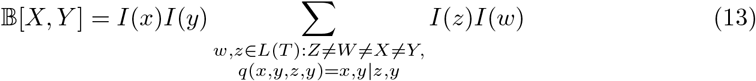

where the importance values of singletons are set to 1 so we know that *I*(*x*) = *I*(*y*) = 1. Note that all of the importance values are set to 1 in the unnormalized case.

To compute the normalized version of 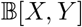 using the previous algorithm, we set *c_D_*[*v*] to be the sum of the importance values of the leaves below *v* that are labeled by *D* (i.e., *c_D_*[*v*] ∑_*m*∈*L*(*v*), *M*=*D*_ *I*(*m*) where *L*(*v*) denotes the set of leaves below *v*). The proof of correctness follows from Lemma 1, in which we show that the total weight of selecting two uniquely labeled leaves below vertex *u* equals *g*_0_[*u*]. Intuitively, this is because all other quantities (*p*, 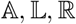) are computed from *g*_0_[*u*].

#### Lemma 1.

*The total weight of all taxon pairs in the subtree rooted at internal vertex u*

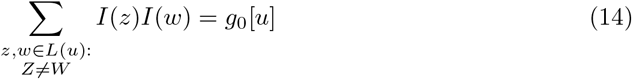

*where L*(*u*) *is the set of leaves below vertex u*.

See Supplementary Materials for proof.

Lastly, we need to compute the good edges 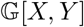, which is the total weight of quartets in which *X, Y* are not siblings. This can be done in constant time, following Lemma 2.

#### Lemma 2.

*Let T be a multi-labeled gene tree, and let X, Y be singletons. Then*,

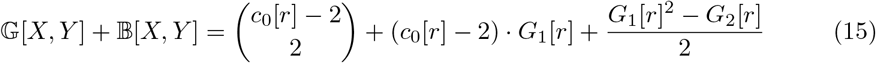

*where r is the root vertex of T*.

See Supplementary Materials for proof.

This concludes our treatment of case 1, in which *X, Y* are both singletons. In order to compute all entries of 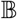 and 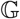, we also need to consider the other two cases. In case 2, *X* is a singleton and *Y* is an artificial taxon (or vice versa), and in case 3, both *X* and *Y* are artificial taxa. These cases are more complicated because the naive approach would consider all paths in the tree between a leaf labeled *X* and a leaf labeled *Y*, which is not efficient. The algorithms and proofs for these cases are provided in the Supplementary Materials.

## Software and Data Availability

TREE-QMC is available on Github: https://github.com/molloy-lab/TREE-QMC. The scripts used to run methods and analyze the results are also available on Github: https://github.com/molloy-lab/tree-qmc-study. The data (including true and estimated gene trees as well as true and estimated species trees) are available on Dryad: https://doi.org/10.5061/dryad.m0cfxpp6g.

## Data Access

This research did not generate new data.

## Acknowledgments

This research was funded by the State of Maryland.

