## Supplementary Materials for "TREE-QMC: Improving quartet graph construction for scalable and accurate species tree estimation from gene trees"

December 30, 2022

#### Contents

**List of Tables** **2**

**List of Figures** **2**

**1 Divide-and-Conquer Framework** **3**

**2 Details of Experimental Study** **4**

**References** **34**

### 1 Divide-and-Conquer Framework

Figure S1: **Divide-and-conquer framework utilized by wQMC, TREE-QMC, and wQFM.** Initially, the input data (either gene trees or weighted quartets) are on species set $\mathcal{S} = \{0, 1, \ldots, 10\}$. At each step in the divide phase, we identify a bipartition and recurse on the implied subproblems. For problem $P_0$, we identify bipartition $\mathcal{E}|\mathcal{F}$, where $\mathcal{E} = \{0, 2, 4, 8, 9\}$ and $\mathcal{F} = \{1, 3, 5, 6, 7, 10\}$. We then introduce artifical taxa $A$ (representing the taxa in $\mathcal{F}$) and $a$ (representing the taxa in $\mathcal{E}$) and recurse on species sets $\{A\} \cup \mathcal{E}$ (subproblem $P_1$) and $\{a\} \cup \mathcal{F}$ (subproblem $P_2$), updating the input data (either gene trees or weighted quartets) accordingly. We continue in this fashion until there are fewer than four taxa (e.g., problem $P_7$) in which case we return the only possible tree (e.g., tree $T_7$). At each step in the conquer phase, we connect pairs of trees using the artifical taxa. On the final conquer step, we connect trees $T_1$ and $T_2$ at artificial taxa $A$ and $a$, producing a tree on species set $\mathcal{S}$.

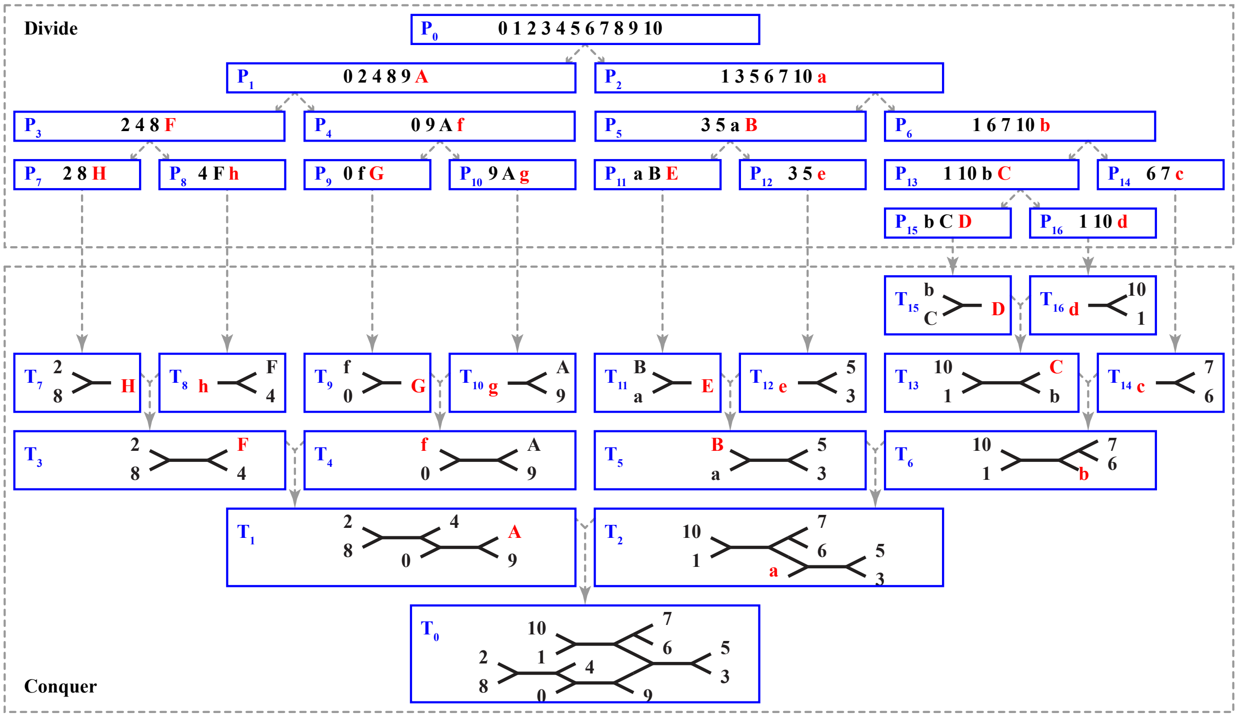

**Subproblems and Artificial Taxa.** At each step in the divide phase, wQMC finds a bipartition on species set $\mathcal{X}$ using the input quartets $\mathcal{Q}_\mathcal{X}$. The the quartets are then updated based on the implied subproblems (Avni et al. 2014). Given bipartition $\mathcal{E}|\mathcal{F}$, a quartet $q = A, B|C, D \in \mathcal{Q}_\mathcal{X}$ is assigned to one of four cases:

1. *satisfied* if $A, B \in \mathcal{E}$ and $C, D \in \mathcal{F}$ (or vice versa),
2. *violated* if
   - $A, C \in \mathcal{E}$ and $B, D \in \mathcal{F}$ (or vice versa) or
   - $A, D \in \mathcal{E}$ and $B, C \in \mathcal{F}$ (or vice versa),
3. *untouched* if $|\{A, B, C, D\} \cap \mathcal{E}| = 4$ or $|\{A, B, C, D\} \cap \mathcal{F}| = 4$, and
4. *deferred* if $|\{A, B, C, D\} \cap \mathcal{E}| = 3$ or $|\{A, B, C, D\} \cap \mathcal{F}| = 3$.

If the quartet is satisfied or violated, it is discarded because it will be satisfied or violated regardless of the bipartitions produced at future steps in the divide phase of the algorithm. If the quartet is untoched or deferred, it is assigned to one of the two subproblems: $\mathcal{Q}_{\mathcal{E}+}$ or $\mathcal{Q}_{\mathcal{F}+}$, where $\mathcal{Q}_{\mathcal{E}+}$ denotes the set of quartets formed by adding the untouched and deferred quartets with 4 or 3 leaves labeled by elements of $\mathcal{E}$ (and similarly for $\mathcal{Q}_{\mathcal{F}+}$). To add a deferred quartet to $\mathcal{Q}_{\mathcal{E}+}$, its one leaf labeled by an element of $\mathcal{F}$ is relabeled with artificial taxon $F$. After this relabeling process has been completed for all deferred quartets, there can be multiples of the same quartet (each with its own weight). These multiples are replaced by a single quartet whose weight is the sum.

### 2 Details of Experimental Study

#### 2.1 Replicates Excluded from ASTRAL-II Data

All comparisons between methods are made on the same set of replicate data sets. For analyses of estimated gene trees, we excluded replicate data set from analyses whenever more than half of the 1000 gene trees had the majority of their branches unresolved. Specifically, we excluded

- 1 replicate (# 41) from the 10-taxon model condition
- 2 replicates (# 21, 41) from the 50-taxon model condition
- 2 replicates (# 8, 47) from the 100-taxon model condition
- 3 replicates (# 8, 15, 49) from the 200 taxon model condition with very high ILS and shallow speciation

We also excluded replicates on which ASTRAL-III failed to complete; this occurred only for 3 replicates (# 6, 8, 38) from the 1000-taxon, 1000-gene model condition.

#### 2.2 Properties of Simulated Data

Table S1: **Properties of ASTRAL-II data sets.** ILS for each replicate is quantified as the normalized RF distance between the true species tree and the true gene tree average across all 1000 gene trees. GTEE for each replicate is quantified as the normalized RF distance between the true and estimated gene trees averaged across all 1000 gene trees. AD for each replicate is the normalized RF distance between the true species tree and estimated gene tree averaged across all 1000 gene trees. The values in the table is the average ($\pm$ standard deviation) across all replicates.

| species tree height | speciation | # of taxa | ILS | GTEE | AD |
|---|---|---|---|---|---|
| 0.5X | deep | 200 | $0.68 \pm 0.02$ | $0.44 \pm 0.14$ | $0.74 \pm 0.03$ |
| 0.5X | shallow | 200 | $0.69 \pm 0.02$ | $0.44 \pm 0.12$ | $0.74 \pm 0.03$ |
| 1X | shallow | 10 | $0.17 \pm 0.06$ | $0.19 \pm 0.09$ | $0.28 \pm 0.08$ |
| 1X | shallow | 50 | $0.31 \pm 0.04$ | $0.26 \pm 0.11$ | $0.42 \pm 0.08$ |
| 1X | shallow | 100 | $0.33 \pm 0.02$ | $0.26 \pm 0.09$ | $0.44 \pm 0.05$ |
| 1X | deep | 200 | $0.34 \pm 0.02$ | $0.34 \pm 0.12$ | $0.47 \pm 0.08$ |
| 1X | shallow | 200 | $0.34 \pm 0.02$ | $0.27 \pm 0.12$ | $0.44 \pm 0.07$ |
| 1X | shallow | 500 | $0.34 \pm 0.01$ | $0.28 \pm 0.11$ | $0.45 \pm 0.07$ |
| 1X | shallow | 1000 | $0.35 \pm 0.01$ | $0.30 \pm 0.11$ | $0.47 \pm 0.08$ |
| 5X | deep | 200 | $0.09 \pm 0.01$ | $0.28 \pm 0.11$ | $0.31 \pm 0.10$ |
| 5X | shallow | 200 | $0.21 \pm 0.02$ | $0.21 \pm 0.13$ | $0.33 \pm 0.09$ |

Table S2: **Properties of avian and mammalian simulated data.** Downloaded data sets Mahbub et al. (2021), which did not include true gene trees for the 0.5X and 2X species tree scales for mammalian data set.

| species tree scale | sequence length | ILS | GTEE | AD |
|---|---|---|---|---|
| *Avian (48 taxa, 1000 estimated gene trees)* | | | | |
| 0.5X | 500 | $0.591 \pm 0.002$ | $0.597 \pm 0.094$ | $0.687 \pm 0.001$ |
| 1X | 500 | $0.473 \pm 0.002$ | $0.597 \pm 0.066$ | $0.637 \pm 0.002$ |
| 2X | 500 | $0.354 \pm 0.002$ | $0.619 \pm 0.002$ | $0.597 \pm 0.002$ |
| *Mammalian (37 taxa, 200 estimated gene trees)* | | | | |
| 1X | 250 | $0.330 \pm 0.005$ | $0.427 \pm 0.009$ | $0.538 \pm 0.007$ |
| 1X | 500 | $0.330 \pm 0.005$ | $0.282 \pm 0.058$ | $0.438 \pm 0.006$ |
| 1X | 1000 | $0.330 \pm 0.005$ | $0.161 \pm 0.006$ | $0.380 \pm 0.006$ |
| 1X | 1500 | $0.330 \pm 0.005$ | $0.119 \pm 0.005$ | $0.362 \pm 0.006$ |

#### 2.3 Computational Resources and Empirical Runtime

Running time is reported as the wall-clock time (i.e., the amount of time that elapses between the beginning and end of the computation). All computational experiments were performed on cluster nodes outfitted with 128 GB RAM and 32 cores (2 × 16-Core AMD Opteron(tm) Processor 6378 2400MHz) and running Red Hat Enterprise Linux 7 (x86_64) operating system. There are 12 nodes with these specifications, which can be accessed via a queue that allows exclusive access but limits the allowed memory to 36 GB. Therefore, we ran methods with exclusive access to the node with access to 36GB of memory, assigning all analyses of the same replicate data set to the same node. All five methods we evaluated are single-threaded.

#### 2.4 Species Tree Estimation Commands

##### 2.4.1 ASTRAL, FASTRAL, and TREE-QMC.

ASTRAL version 5.7.7 (Zhang et al. 2018) was run with the following command:

```
java -Xmx36G -D"java.library.path=<path to ASTRAL>/lib" -jar <path to ASTRAL>/astral.5.7.7.jar \
     -t0 -i [input gene trees] -o [output species tree] &> [output log file]
```

Note that `-t0` means that branch length and support estimation are not performed.
FASTRAL (Dibaeinia et al. 2021) was run with the following command:

```
fastral --ns 1,10,20,20 --nt 1000,500,250,100 --k 1000 \
        --it [input gene trees] --os [output directory]/samples \
        --aggregate [output directory]/[output constraints] \
        --o [output directory]/[output species tree] \
        --time [output directory]/[output timing file] \
        --path_ASTRAL [ASTRAL path] \
        --path_ASTRID [ASTRID path] &> [output log file]
```

Note that `-nt 250,125,63,25 -k 250` where used for data sets with 250 genes. We ran FASTRAL with the version of ASTRAL and ASTRID distributed with the code; however, we modified the command to call ASTRAL so that it could use 36 GB of RAM and so that it did NOT compute branch support for the final tree.

TREE-QMC version 1.0.0 was run with the following command:

```
TREEQMC -n 0 -i [input gene trees] -o [output species tree] &> [output log file]
```

Note that `-n 1` and `-n 2` where also run and in this case the input gene trees were the refined trees from running TREE-QMC with `-n 0`. This is so the only difference between run of TREE-QMC is the normalization scheme (and not the refinement of polytomies in the input gene trees).

##### 2.4.2 wQMC and wQFM.

Unlike the previous methods, wQMC and wQFM take weighted quartets as input. We used the scripts provided with wQFM (`https://github.com/Mahim1997/wQFM-2020`) to extract weighted quartets from the gene trees. Specifically, we used the command:

```
quartet-controller.sh [input gene trees] [output weighted quartets] &> [output log file]
```

which calls by `triplets.soda2103` executable from (). We then selected the binary quartets

```
grep "),(" [input weighted quartets] >> [output weighted quartets for wQFM]
```

and re-formatted them for wQMC:

```
sed 's/),(/|/g' [input weighted quartets for wQFM] | \
     sed 's/(//g' | sed 's/)//g' | sed 's/; /:/g' \
     > [output weighted quartets for wQMC]
```

wQFM version 1.3 (Mahbub et al. 2021) was run with the following command:

```
java -Xmx36G -jar [path to wQFM]/wQFM-v1.3.jar \
     -i [input weighted quartets] -o [output species tree] &> [output log file]
```

wQMC version 3.0 (Avni et al. 2014) was run with the following command:

```
max-cut-tree qrtt=[input weighted quartets] \
             weights=on otre=[output species tree] &> [output log file]
```

##### 2.5 Quartet Score and Branch Support Estimation Commands

Quartet score (and branch support) was computed for estimated species trees with the following command:

```
java -Xmx36G -D"java.library.path=<path to ASTRAL>/lib" -jar <path to ASTRAL>/astral.5.7.7.jar \
     -t2 -q [input species tree] -i [input gene trees] \
     -o [output scored species tree] &> [output log file]
```

Table S3: **Runtimes for ASTRAL-II data (1000 estimated gene trees).** Runtime (minutes) is reported for all methods (note: a runtime of zero minutes means it rounded down). Values shown are averages (± standard deviations) across across replicate data sets on which all methods completed (note: ASTRAL-III did not complete on 3 replicates with 1000 taxa and 1000 genes). Lowest median values are in bold. The value in parentheses for wQMC and wQFM is the fraction of the runtime spent weighting quartets to give to these methods as input.

| Species tree height | # of taxa | spec. | ASTRAL-III | FASTRAL | TREE-QMC n0 | TREE-QMC n1 | TREE-QMC n2 | wQMC | wQFM |
|---|---|---|---|---|---|---|---|---|---|
| *1000 estimated gene trees* | | | | | | | | | |
| 0.5X | 200 | deep | 77.4 ± 27.9 | **1.6 ± 0.4** | 2.7 ± 0.5 | 3.0 ± 0.6 | 3.0 ± 0.6 | NA | NA |
| 0.5X | 200 | shallow | 72.7 ± 29.0 | **1.5 ± 0.3** | 2.6 ± 0.5 | 2.8 ± 0.5 | 2.8 ± 0.3 | NA | NA |
| 2X | 10 | shallow | **0.0 ± 0.0** | 0.1 ± 0.0 | **0.0 ± 0.0** | **0.0 ± 0.0** | **0.0 ± 0.0** | 0.2 ± 0.0 (0.99) | 0.2 ± 0.0 (0.94) |
| 2X | 50 | shallow | 0.6 ± 0.6 | 0.3 ± 0.0 | **0.2 ± 0.0** | **0.2 ± 0.0** | **0.2 ± 0.0** | 15.8 ± 3.0 (1.00) | 16.8 ± 3.0 (0.94) |
| 2X | 100 | shallow | 3.0 ± 2.4 | 0.7 ± 0.1 | **0.6 ± 0.1** | **0.6 ± 0.1** | **0.6 ± 0.1** | 256.8 ± 34.7 (1.00) | 290.2 ± 38.4 (0.88) |
| 2X | 200 | deep | 17.1 ± 11.4 | **1.2 ± 0.1** | 2.4 ± 0.1 | 2.5 ± 0.2 | 2.5 ± 0.2 | NA | NA |
| 2X | 200 | shallow | 12.0 ± 9.5 | **1.2 ± 0.2** | 2.4 ± 0.4 | 2.5 ± 0.4 | 2.5 ± 0.4 | NA | NA |
| 2X | 500 | shallow | 72.1 ± 46.2 | **5.6 ± 1.1** | 14.8 ± 2.7 | 15.8 ± 3.2 | 15.5 ± 2.5 | NA | NA |
| 2X | 1000 | shallow | 317.8 ± 198.9 | **32.4 ± 7.7** | 59.2 ± 11.6 | 62.0 ± 9.2 | 63.5 ± 13.2 | NA | NA |
| 5X | 200 | deep | 6.6 ± 7.7 | **1.2 ± 0.0** | 2.2 ± 0.1 | 2.3 ± 0.1 | 2.3 ± 0.1 | NA | NA |
| 5X | 200 | shallow | 4.6 ± 7.9 | **1.1 ± 0.0** | 2.2 ± 0.0 | 2.3 ± 0.0 | 2.3 ± 0.0 | NA | NA |
| *250 estimated gene trees* | | | | | | | | | |
| 0.5X | 200 | deep | 11.3 ± 4.8 | 0.8 ± 0.2 | **0.7 ± 0.1** | **0.7 ± 0.1** | **0.7 ± 0.2** | NA | NA |
| 0.5X | 200 | shallow | 9.6 ± 3.4 | 0.7 ± 0.1 | **0.6 ± 0.0** | 0.7 ± 0.0 | 0.7 ± 0.0 | NA | NA |
| 2X | 10 | shallow | **0.0 ± 0.0** | 0.1 ± 0.0 | **0.0 ± 0.0** | **0.0 ± 0.0** | **0.0 ± 0.0** | 0.1 ± 0.0 (0.98) | 0.1 ± 0.0 (0.83) |
| 2X | 50 | shallow | 0.1 ± 0.0 | 0.1 ± 0.0 | **0.0 ± 0.0** | **0.0 ± 0.0** | **0.0 ± 0.0** | 4.1 ± 0.8 (0.99) | 5.1 ± 1.0 (0.80) |
| 2X | 100 | shallow | 0.3 ± 0.2 | **0.2 ± 0.0** | **0.2 ± 0.0** | **0.2 ± 0.0** | **0.2 ± 0.0** | 65.4 ± 8.2 (1.00) | 96.4 ± 12.5 (0.68) |
| 2X | 200 | deep | 1.7 ± 1.2 | **0.5 ± 0.0** | 0.6 ± 0.0 | 0.6 ± 0.0 | 0.6 ± 0.0 | NA | NA |
| 2X | 200 | shallow | 1.1 ± 0.7 | **0.5 ± 0.1** | 0.6 ± 0.1 | 0.6 ± 0.1 | 0.6 ± 0.2 | NA | NA |
| 2X | 500 | shallow | 7.1 ± 4.7 | **3.0 ± 0.8** | 3.7 ± 0.5 | 3.9 ± 0.7 | 3.9 ± 0.8 | NA | NA |
| 2X | 1000 | shallow | 45.9 ± 50.4 | 22.4 ± 9.4 | **15.2 ± 2.5** | 15.8 ± 2.2 | 15.5 ± 1.5 | NA | NA |
| 5X | 200 | deep | 0.6 ± 0.6 | **0.4 ± 0.0** | 0.6 ± 0.0 | 0.6 ± 0.0 | 0.6 ± 0.0 | NA | NA |
| 5X | 200 | shallow | 0.5 ± 0.6 | **0.4 ± 0.0** | 0.6 ± 0.0 | 0.6 ± 0.0 | 0.6 ± 0.0 | NA | NA |

### 3 Additional Results on Simulated Data Sets

#### 3.1 Number of taxa

Table S4: **Testing for differences between TREE-QMC-n2 and other methods on ASTRAL-II data sets with varying numbers of taxa and 1000 estimated gene trees (Figure 2).** All data sets were simulated with 1X species tree height and shallow speciation. The table shows the number of replicates for which species tree estimation error was better for TREE-QMC-n2, better for the other method, or a tie as well as other metrics. The $\Delta$ false negative (FN) error is the difference in the number of false negative branches between the other method and TREE-QMC-n2, averaged ($\pm$ standard deviation) across all replicates (including ties). Positive values indicate that TREE-QMC-n2 is better on average, whereas negative values indicate that the other method is better on average. Differences between methods were evaluated using two-sided Wilcoxon signed-rank tests, after removing tied values (McGee 2018) (note: the test was performed only if more than than 25 replicates were not ties). The symbol *, *, *** indicate significance at $p < 0.05$, 0.005, and 0.0005, respectively. Note that $p < 0.05/12 = 0.0041\overline{6}$ survives Bonferroni correction for the 12 comparisons made in this table (MC).

| # of taxa | Other method | # of replicates TQMC-n2 | OTHER | TIE | $\Delta$ FN error | p-value | | MC |
|---|---|---|---|---|---|---|---|---|
| 10 | FASTRAL | 0 | 0 | 49 | 0.00 ± 0.00 | NA | | |
| 10 | ASTRAL3 | 0 | 0 | 49 | 0.00 ± 0.00 | NA | | |
| 50 | FASTRAL | 5 | 3 | 40 | 0.04 ± 0.41 | NA | | |
| 50 | ASTRAL3 | 5 | 3 | 40 | 0.04 ± 0.41 | NA | | |
| 100 | FASTRAL | 9 | 8 | 31 | 0.04 ± 0.92 | NA | | |
| 100 | ASTRAL3 | 10 | 8 | 30 | -0.08 ± 1.49 | NA | | |
| 200 | FASTRAL | 20 | 6 | 24 | 0.44 ± 1.09 | 0.006217 | * | N |
| 200 | ASTRAL3 | 20 | 6 | 24 | 0.48 ± 1.13 | 0.004217 | ** | Y |
| 500 | FASTRAL | 34 | 9 | 7 | 2.40 ± 3.46 | 0.000009 | *** | Y |
| 500 | ASTRAL3 | 33 | 8 | 9 | 2.70 ± 3.89 | 0.000013 | *** | Y |
| 1000 | FASTRAL | 32 | 10 | 5 | 3.94 ± 6.96 | 0.000123 | *** | Y |
| 1000 | ASTRAL3 | 35 | 8 | 4 | 4.64 ± 7.84 | 0.000008 | *** | Y |

#### 3.2 Species tree scale/height and thus ILS level

Table S5: **Testing for differences between TREE-QMC-n2 and other methods on ASTRAL-II data sets with varying ILS levels and 1000 estimated gene trees (Figure 3)**. All data sets have 200 taxa. Note: rows with parentheses are duplicated from Table S4.

| species tree height | speciation | Other method | # of replicates TQMC-n2 | OTHER | TIE | Δ FN error | p-value | | MC |
|---|---|---|---|---|---|---|---|---|---|
| 0.25X | deep | FASTRAL | 22 | 21 | 7 | -0.24 ± 3.46 | 0.70 | | |
| 0.25X | deep | ASTRAL3 | 29 | 18 | 3 | 0.5 ± 3.52 | 0.10 | | |
| 0.25X | shallow | FASTRAL | 23 | 16 | 8 | 0.47 ± 2.24 | 0.16 | | |
| 0.25X | shallow | ASTRAL3 | 23 | 15 | 9 | 0.77 ± 2.35 | 0.04414 | * | N |
| 1X | deep | FASTRAL | 20 | 9 | 21 | 0.70 ± 2.15 | 0.04209 | * | N |
| 1X | deep | ASTRAL3 | 21 | 7 | 22 | 1.00 ± 2.25 | 0.00367 | ** | Y |
| (1X) | (shallow) | FASTRAL | 20 | 6 | 24 | 0.44 ± 1.09 | 0.00622 | * | N |
| (1X) | (shallow) | ASTRAL3 | 20 | 6 | 24 | 0.48 ± 1.13 | 0.00422 | ** | N |
| 5X | deep | FASTRAL | 27 | 6 | 17 | 1.10 ± 1.88 | 0.00032 | *** | Y |
| 5X | deep | ASTRAL3 | 25 | 8 | 17 | 0.90 ± 2.26 | 0.00382 | ** | Y |
| 5X | shallow | FASTRAL | 13 | 8 | 29 | 0.44 ± 1.39 | NA | | |
| 5X | shallow | ASTRAL3 | 13 | 9 | 28 | 0.42 ± 1.42 | NA | | |

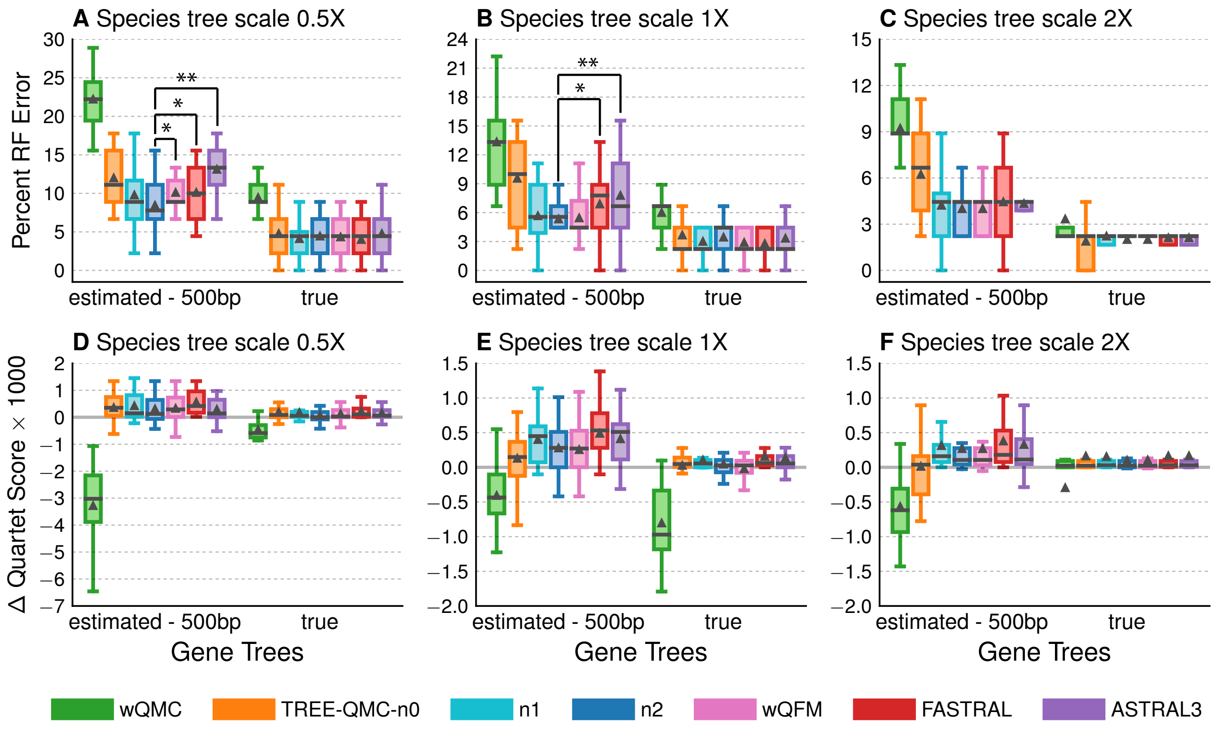

Figure S2: Reproduction of Figure 4 in main text with different y-axis.

Table S6: **Testing for differences between TREE-QMC-n2 and other methods on avian simulated data sets with species tree scales and thus ILS levels (Figure 4).** All data sets had 1000 gene trees. Differences between methods were evaluated using two-sided Wilcoxon signed-rank tests, after removing tied values (McGee 2018) (note: the test was only performed if more than than 10 replicates were not ties). The symbol *, **, and *** indicate significance at $p < 0.05$, 0.005, and 0.0005. Note: $p < 0.05/18 = 0.00278$ would be significant after Bonferroni correction for the 18 comparisons made in this table (MC).

| Gene trees | Other method | # of replicates TQMC-n2 | OTHER | TIE | Δ FN error | p-value | | MC |
|---|---|---|---|---|---|---|---|---|
| *Figure 4A – 0.5X species tree scale* | | | | | | | | |
| estimated | wQFM | 11 | 2 | 7 | 0.75 ± 1.12 | 0.011006 | * | N |
| estimated | FASTRAL | 8 | 3 | 9 | 0.75 ± 1.48 | 0.0461 | * | N |
| estimated | ASTRAL3 | 18 | 2 | 0 | 2.10 ± 1.86 | 0.00101 | ** | Y |
| true | wQFM | 5 | 7 | 8 | -0.05 ± 1.340 | 0.94 | | |
| true | FASTRAL | 5 | 8 | 7 | -0.20 ± 1.24 | 0.56 | | |
| true | ASTRAL3 | 9 | 7 | 4 | 0.15 ± 1.42 | 0.56 | | |
| *Figure 4B – 1X species tree scale* | | | | | | | | |
| estimated | wQFM | 5 | 4 | 11 | 0.05 ± 0.69 | NA | | |
| estimated | FASTRAL | 9 | 3 | 8 | 0.70 ± 1.34 | 0.0333 | * | N |
| estimated | ASTRAL3 | 12 | 1 | 7 | 1.1 ± 1.21 | 0.00293 | ** | N |
| true | wQFM | 2 | 6 | 12 | -0.25 ± 0.72 | NA | | |
| true | FASTRAL | 2 | 7 | 11 | -0.30 ± 0.92 | NA | | |
| true | ASTRAL3 | 4 | 5 | 11 | -0.05 ± 0.89 | NA | | |
| *Figure 4C – 2X species tree scale* | | | | | | | | |
| estimated | wQFM | 4 | 5 | 11 | 0.00 ± 0.80 | NA | | |
| estimated | FASTRAL | 7 | 5 | 8 | 0.20 ± 0.95 | 0.36 | | |
| estimated | ASTRAL3 | 7 | 5 | 8 | 0.15 ± 0.88 | 0.46 | | |
| true | wQFM | 0 | 0 | 20 | 0.00 ± 0.00 | NA | | |
| true | FASTRAL | 2 | 1 | 17 | 0.05 ± 0.39 | NA | | |
| true | ASTRAL3 | 2 | 1 | 17 | 0.05 ± 0.39 | NA | | |

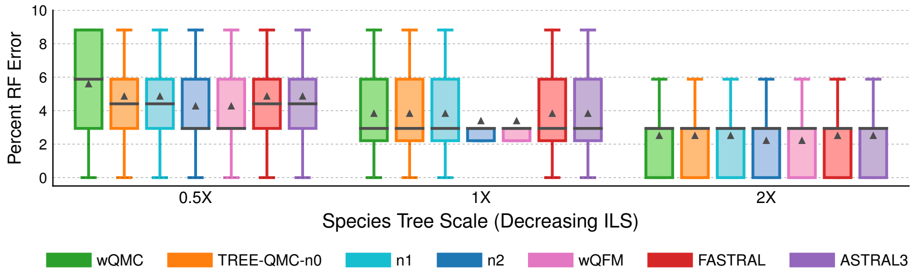

Figure S3: **Species tree error and runtime for mammalian simulated data set with varying ILS levels.** All data sets have 200 estimated gene trees (sequence length: 500). Percent species tree error across replicates is shown using box plots (bars represent medians; triangles represent means; some outliers are not shown).

Table S7: **Testing for differences between TREE-QMC-n2 and other methods on mammalian simulated data sets with varying species tree scales and thus ILS levels (Figure S3).** All data sets had 200 estimated gene trees (sequence length: 500).

| Species tree scale | Other method | # of replicates TQMC-n2 | OTHER | TIE | $\Delta$ FN error | p-value |
|---|---|---|---|---|---|---|
| 0.5X | wQFM | 0 | 0 | 20 | $0.00 \pm 0.00$ | NA |
| 0.5X | FASTRAL | 5 | 2 | 13 | $0.20 \pm 0.70$ | NA |
| 0.5X | ASTRAL3 | 5 | 2 | 13 | $0.20 \pm 0.70$ | NA |
| 1X | wQFM | 0 | 0 | 20 | $0.00 \pm 0.00$ | NA |
| 1X | FASTRAL | 5 | 2 | 13 | $0.15 \pm 0.59$ | NA |
| 1X | ASTRAL3 | 5 | 2 | 13 | $0.15 \pm 0.59$ | NA |
| 2X | wQFM | 0 | 0 | 20 | $0.00 \pm 0.00$ | NA |
| 1X | FASTRAL | 3 | 1 | 16 | $0.10 \pm 0.45$ | NA |
| 2X | ASTRAL3 | 3 | 1 | 16 | $0.10 \pm 0.45$ | NA |

#### 3.3 Sequence length and thus GTEE level

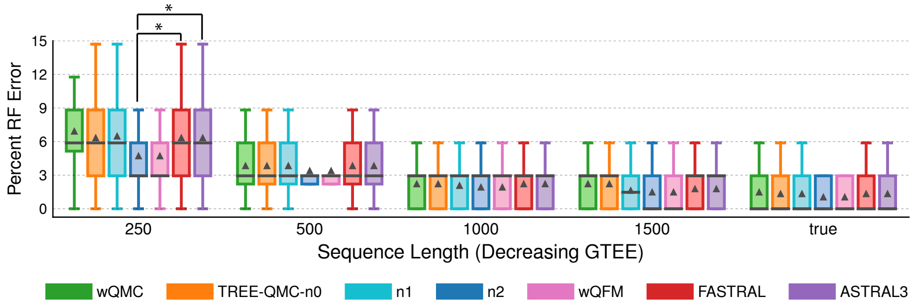

Figure S4: **Species tree error and runtime for mammalian simulated data set with varying sequence lengths and GTEE levels.** All data sets were simulated from unscaled species trees (1X) and have 200 gene trees (note: results for sequence length 500 are duplicated from Figure S3). Percent species tree error across replicates is shown using box plots (bars represent medians; triangles represent means; some outliers are not shown).

Table S8: **Testing for differences between TREE-QMC-n2 and other methods on mammalian simulated data sets with varying sequence lengths and thus gene tree estimation error (Figure S4).** All data sets were simulated with from unscaled species trees (1X) and have 200 gene trees. Note: rows with parentheses are duplicated from Table S7.

| Sequence length | Other method | # of replicates TQMC-n2 | OTHER | TIE | Δ FN error | p-value | |
|---|---|---|---|---|---|---|---|
| 250 | wQFM | 0 | 0 | 20 | 0.00 ± 0.00 | NA | |
| 250 | FASTRAL | 11 | 4 | 5 | 0.55 ± 1.10 | 0.04 | * |
| 250 | ASTRAL3 | 11 | 4 | 5 | 0.55 ± 1.10 | 0.04 | * |
| (500) | wQFM | 0 | 0 | 20 | 0.00 ± 0.00 | NA | |
| (500) | FASTRAL | 5 | 2 | 13 | 0.15 ± 0.59 | NA | |
| (500) | ASTRAL3 | 5 | 2 | 13 | 0.15 ± 0.59 | NA | |
| 1000 | wQFM | 0 | 0 | 20 | 0.00 ± 0.00 | NA | |
| 1000 | FASTRAL | 2 | 0 | 18 | 0.10 ± 0.31 | NA | |
| 1000 | ASTRAL3 | 2 | 0 | 18 | 0.10 ± 0.31 | NA | |
| 1500 | wQFM | 0 | 0 | 20 | 0.00 ± 0.00 | NA | |
| 1500 | FASTRAL | 3 | 1 | 16 | 0.10 ± 0.45 | NA | |
| 1500 | ASTRAL3 | 3 | 1 | 16 | 0.10 ± 0.45 | NA | |
| true | wQFM | 0 | 0 | 20 | 0.00 ± 0.00 | NA | |
| true | FASTRAL | 3 | 1 | 16 | 0.10 ± 0.45 | NA | |
| true | ASTRAL3 | 3 | 1 | 16 | 0.10 ± 0.45 | NA | |

#### 3.4 Number of Gene Trees

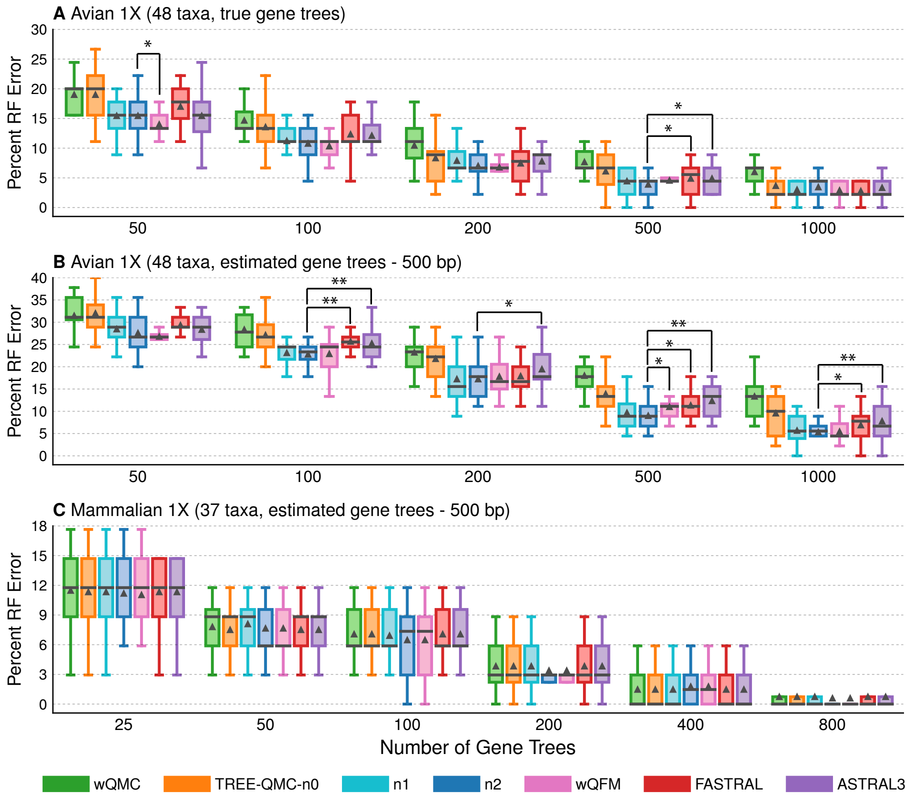

Figure S5: **Species tree error and runtime for avian and mammalian simulated data sets.** Bars represent medians, and triangles represent means (outliers are not shown). The symbols *, **, and *** indicate significance at $p < 0.05$, 0.005, and 0.0005, respectively (see Tables S9 and S10 for details). All data sets have unscaled species trees (1X). Note: the results for avian simulated data sets with 1000 true gene trees are duplicated from Figure 4A–C, and the results for mammalian simulated data sets with 200 estimated gene trees (sequence length: 500) are duplicated from Figures S3 and S4.

Table S9: **Testing for differences between TREE-QMC-n2 and other methods on avian simulated data sets with varying numbers of gene trees (Figure S5).** Note: rows with parentheses are duplicated from Table S6.

| # of genes | Other method | # of replicates TQMC-n2 | OTHER | TIE | Δ FN error | p-value | |
|---|---|---|---|---|---|---|---|
| *Figure S5A – Avian 1X with true gene trees* | | | | | | | |
| 50 | wQFM | 2 | 10 | 8 | -0.65 ± 1.18 | 0.03714 | * |
| 50 | FASTRAL | 11 | 5 | 4 | 0.70 ± 1.56 | 0.07448 | |
| 50 | ASTRAL3 | 8 | 8 | 4 | 0.00 ± 1.41 | 1.00 | |
| 100 | wQFM | 7 | 6 | 7 | -0.20 ± 1.24 | 0.47 | |
| 100 | FASTRAL | 9 | 3 | 8 | 0.70 ± 1.56 | 0.07745 | |
| 100 | ASTRAL3 | 9 | 4 | 7 | 0.60 ± 1.31 | 0.06917 | |
| 200 | wQFM | 2 | 5 | 13 | -0.10 ± 0.97 | NA | |
| 200 | FASTRAL | 8 | 5 | 7 | 0.20 ± 1.20 | 0.49 | |
| 200 | ASTRAL3 | 9 | 4 | 7 | 0.35 ± 1.23 | 0.25 | |
| 500 | wQFM | 5 | 0 | 15 | 0.30 ± 0.57 | NA | |
| 500 | FASTRAL | 10 | 3 | 7 | 0.45 ± 0.89 | 0.04249 | * |
| 500 | ASTRAL3 | 10 | 3 | 7 | 0.45 ± 0.89 | 0.04249 | * |
| (1000) | wQFM | 2 | 6 | 12 | -0.25 ± 0.72 | NA | |
| (1000) | FASTRAL | 2 | 7 | 11 | -0.30 ± 0.92 | NA | |
| (1000) | ASTRAL3 | 4 | 5 | 11 | -0.05 ± 0.89 | NA | |
| *Figure S5B – Avian 1X with estimated gene trees* | | | | | | | |
| 50 | wQFM | 3 | 8 | 9 | -0.35 ± 1.66 | 0.47 | |
| 50 | FASTRAL | 11 | 5 | 4 | 0.80 ± 2.40 | 0.16 | |
| 50 | ASTRAL3 | 8 | 6 | 6 | 0.35 ± 2.48 | 0.51 | |
| 100 | wQFM | 7 | 8 | 5 | 0.05 ± 1.67 | 0.93 | |
| 100 | FASTRAL | 13 | 1 | 6 | 1.30 ± 1.49 | 0.00202 | ** |
| 100 | ASTRAL3 | 13 | 2 | 5 | 1.10 ± 1.37 | 0.00276 | ** |
| 200 | wQFM | 8 | 5 | 7 | 0.25 ± 1.41 | 0.39 | |
| 200 | FASTRAL | 9 | 6 | 5 | 0.30 ± 1.53 | 0.35 | |
| 200 | ASTRAL3 | 11 | 4 | 5 | 1.00 ± 1.75 | 0.02670 | * |
| 500 | wQFM | 11 | 2 | 7 | 0.90 ± 1.45 | 0.01932 | * |
| 500 | FASTRAL | 11 | 3 | 6 | 1.05 ± 1.88 | 0.02633 | * |
| 500 | ASTRAL3 | 15 | 2 | 3 | 1.50 ± 1.50 | 0.00112 | ** |
| (1000) | wQFM | 5 | 4 | 11 | 0.05 ± 0.69 | NA | |
| (1000) | FASTRAL | 9 | 3 | 8 | 0.70 ± 1.34 | 0.03329 | * |
| (1000) | ASTRAL3 | 12 | 1 | 7 | 1.10 ± 1.21 | 0.00293 | ** |

Table S10: **Testing for differences between TREE-QMC-n2 and other methods on mammalian simulated data sets with varying numbers of estimated gene trees (Figure S5).** Note: rows with parentheses are duplicated from Tables S7 and S8.

| # of genes | Other method | # of replicates TQMC-n2 | OTHER | TIE | Δ FN error | p-value |
|---|---|---|---|---|---|---|
| *Figure S5C – Mammalian 1X with estimated gene trees (500 bp)* | | | | | | |
| 25 | wQFM | 0 | 1 | 19 | -0.05 ± 0.22 | NA |
| 25 | FASTRAL | 4 | 5 | 11 | 0.05 ± 0.89 | NA |
| 25 | ASTRAL3 | 4 | 5 | 11 | 0.05 ± 0.89 | NA |
| 50 | wQFM | 0 | 0 | 20 | 0.00 ± 0.00 | NA |
| 50 | FASTRAL | 6 | 5 | 9 | -0.05 ± 0.95 | 0.81 |
| 50 | ASTRAL3 | 6 | 5 | 9 | -0.05 ± 0.95 | 0.81 |
| 100 | wQFM | 0 | 0 | 20 | 0.00 ± 0.00 | NA |
| 100 | FASTRAL | 7 | 3 | 10 | 0.20 ± 0.70 | NA |
| 100 | ASTRAL3 | 7 | 3 | 10 | 0.20 ± 0.70 | NA |
| (200) | wQFM | 0 | 0 | 20 | 0.00 ± 0.00 | NA |
| (200) | FASTRAL | 5 | 2 | 13 | 0.15 ± 0.59 | NA |
| (200) | ASTRAL3 | 5 | 2 | 13 | 0.15 ± 0.59 | NA |
| 400 | wQFM | 0 | 0 | 20 | 0.00 ± 0.00 | NA |
| 400 | FASTRAL | 0 | 2 | 18 | -0.10 ± 0.31 | NA |
| 400 | ASTRAL3 | 0 | 2 | 18 | -0.10 ± 0.31 | NA |
| 800 | wQFM | 0 | 0 | 20 | 0.00 ± 0.00 | NA |
| 800 | FASTRAL | 1 | 0 | 19 | 0.05 ± 0.22 | NA |
| 800 | ASTRAL3 | 1 | 0 | 19 | 0.05 ± 0.22 | NA |

#### 3.5 Additional Results on Biological Data Sets

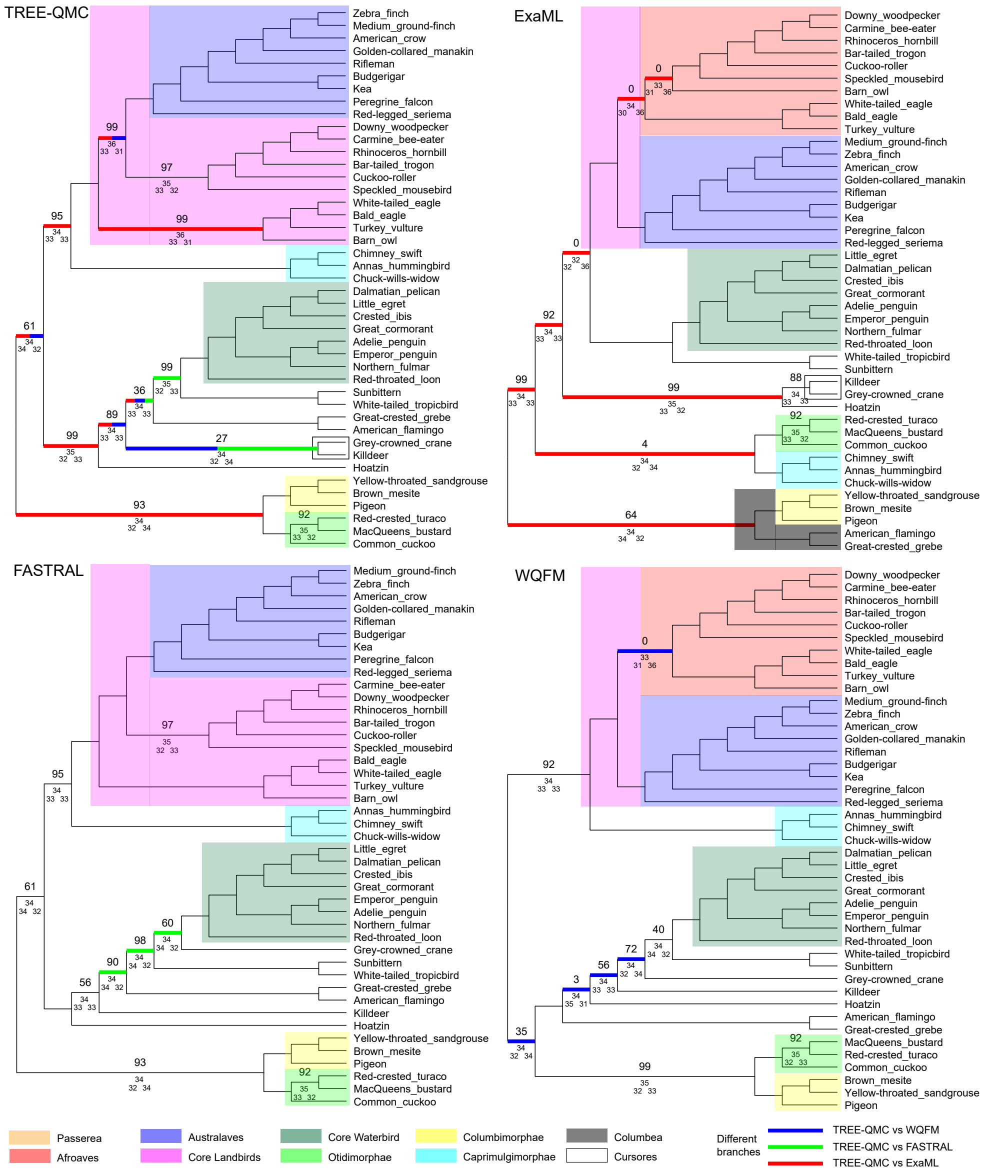

Figure S6: Species trees estimated on TENT data by TREE-QMC-n2, ExaML, FASTRAL, and wQFM. Above the branch, we show support values estimated using ASTRAL's local posterior probability (multiplied by 100). Only support values less than 100 are shown. Below the branch, we show the quartet support (the two values below it correspond to quartet support for the two alternative resolutions of the branch). Importantly, these support calculations are based on estimated gene trees. Taxa outside of Neoaves are not shown as all methods recovered the same topology outside of Neoaves.

### 4 Other Plots for ASTRAL-II Data Sets

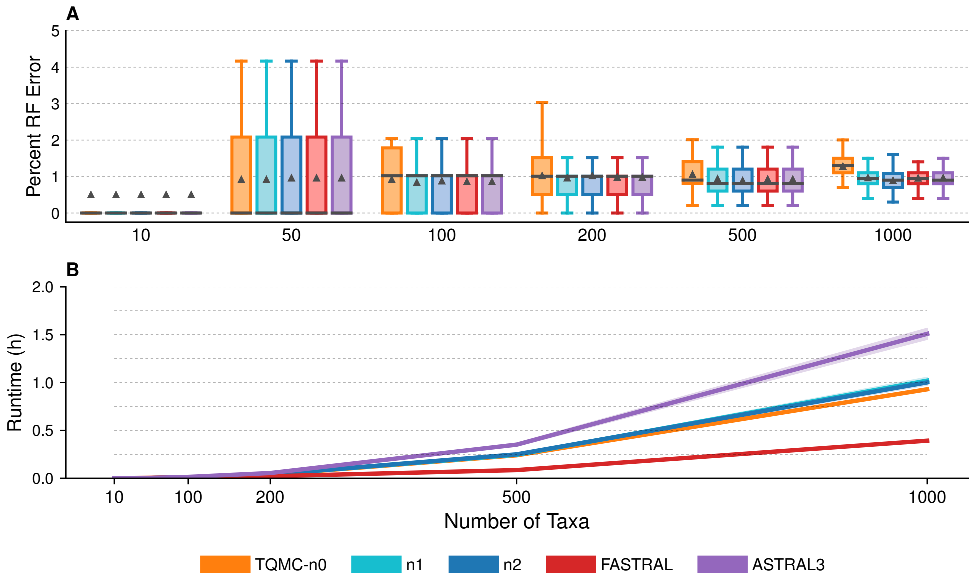

Figure S7: **Species tree error and runtime for ASTRAL-II data sets with varying number of taxa (1000 true gene trees).** (A) Percent species tree error across replicates (bars represent medians; triangles represent means; some outliers are not shown). No statistical tests performed. (B) Mean runtime across replicates (shaded region indicates standard error). All data sets were simulated with 1X species tree height and shallow speciation.

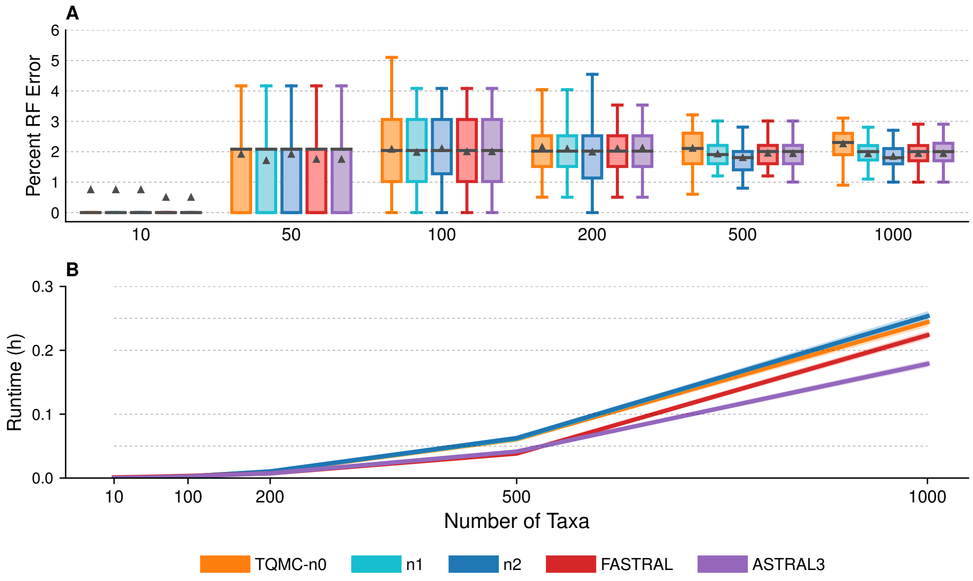

Figure S8: **Species tree error and runtime for ASTRAL-II data sets with varying numbers of taxa (250 true gene trees).** (A) Percent species tree error across replicates (bars represent medians; triangles represent means; some outliers are not shown). No statistical tests performed. (B) Mean runtime across replicates (shaded region indicates standard error). All data sets were simulated with 1X species tree height and shallow speciation.

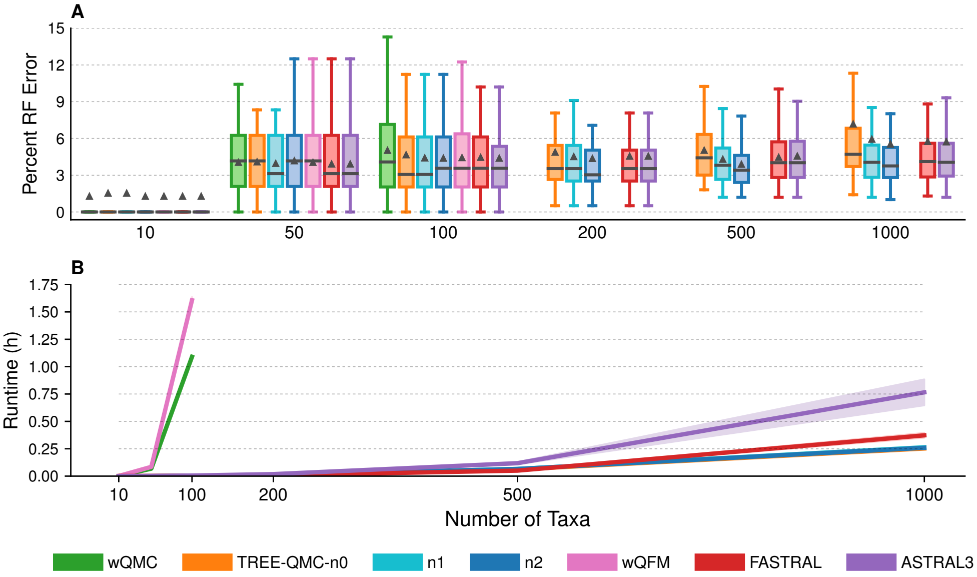

Figure S9: **Species tree error and runtime for ASTRAL-II data sets with varying numbers of taxa (250 estimated gene trees).** (A) Percent species tree error across replicates (bars represent medians; triangles represent means; some outliers are not shown). No statistical tests performed. (B) Mean runtime across replicates (shaded region indicates standard error). All data sets were simulated with 1X species tree height and shallow speciation.

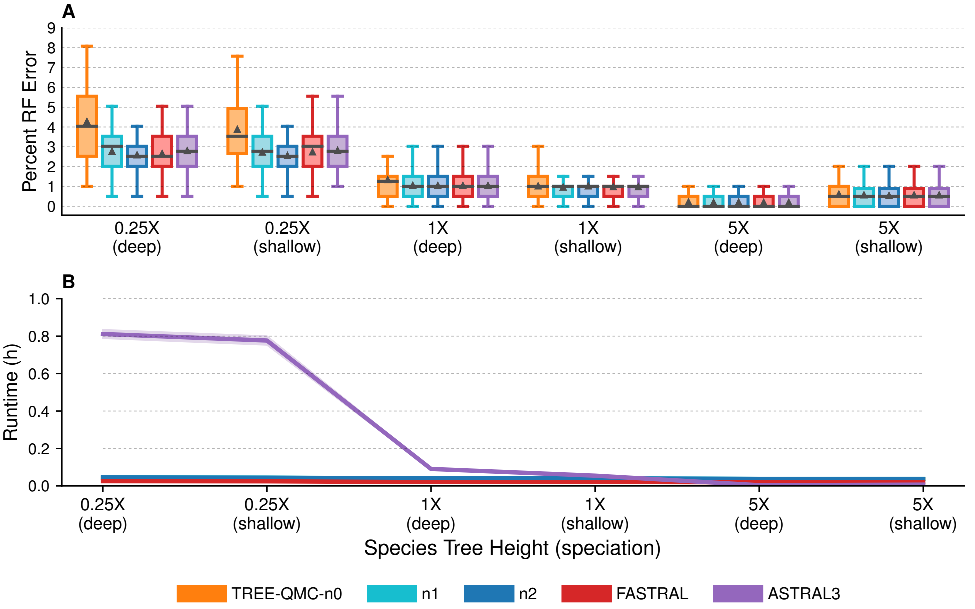

Figure S10: **Species tree error and runtime for ASTRAL-II data sets with varying ILS levels (1000 true gene trees).** (A) Percent species tree error across replicates (bars represent medians; triangles represent means; some outliers are not shown). No statistical tests performed. (B) Mean runtime across replicates (shaded region indicates standard error). All data sets have 200 taxa. One model condition with species tree height 1X and shallow speciation is repeated from Figure S7.

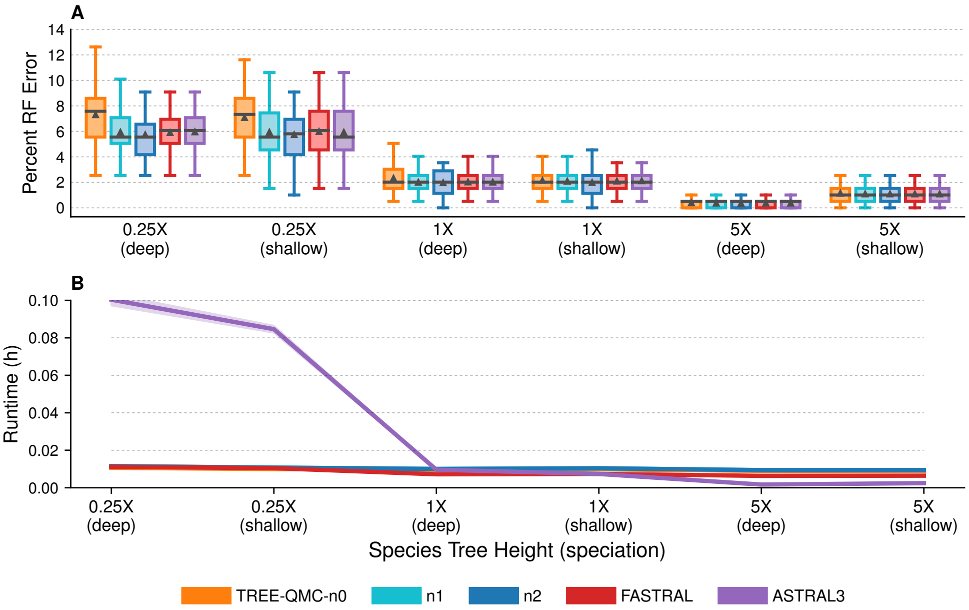

Figure S11: **Species tree error and runtime for ASTRAL-II data sets with varying ILS levels (250 true gene trees).** (A) Percent species tree error across replicates (bars represent medians; triangles represent means; some outliers are not shown). No statistical tests performed. (B) Mean runtime across replicates (shaded region indicates standard error). All data sets have 200 taxa. One model condition with species tree height 1X and shallow speciation is repeated from Figure S8.

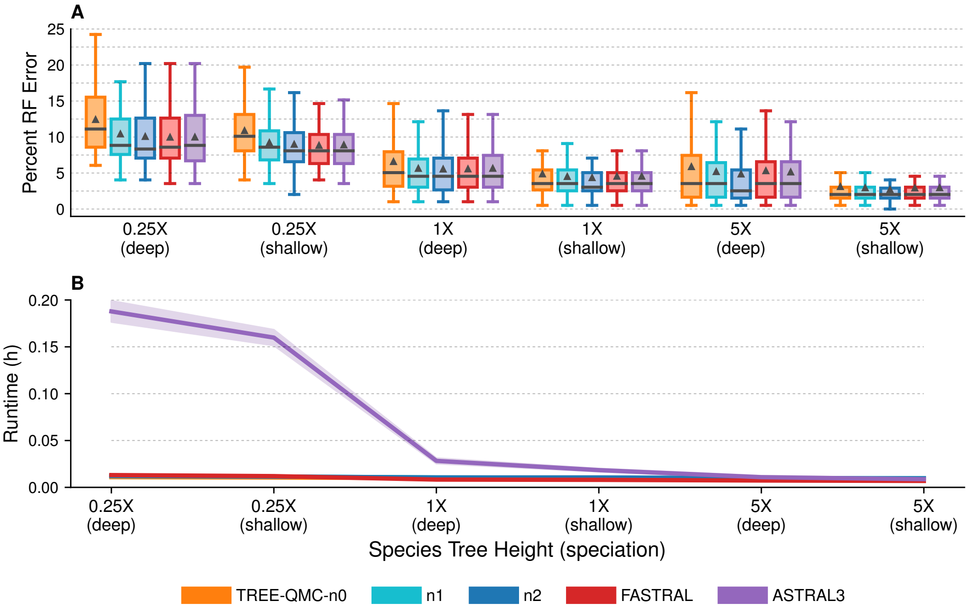

Figure S12: **Species tree error and runtime for ASTRAL-II data sets with varying ILS levels (250 estimated gene trees).** (A) Percent species tree error across replicates (bars represent medians; triangles represent means; some outliers are not shown). No statistical tests performed. (B) Mean runtime across replicates (shaded region indicates standard error). All data sets have 200 taxa. One model condition with species tree height 1X and shallow speciation is repeated from Figure S9.

### 5 TREE-QMC Algorithm

#### 5.1 Quartet Graph Construction Case 1 cont. (Two Singletons)

To compute the normalized version of $\mathbb{B}[X, Y]$ using the previous algorithm, we set $c_D[v]$ to be the sum of the importance values of the leaves below vertex $v$ that are labeled by $D$ (i.e., $c_D[v] = \sum_{m \in L(v), M=D} I(m)$ where $L(v)$ denotes the set of leaves below $v$). The proof of correctness follows from Lemma 1, in which we show that the total weight of selecting two uniquely labeled leaves below vertex $u$ equals $g_0[u]$. All other quantities $(p, \mathbb{A}, \mathbb{L}, \mathbb{R})$ are computed from $g_0[u]$.

**Lemma 1.** *The total weight of all taxon pairs in the subtree rooted at internal vertex $u$*

$$\sum_{\substack{z,w \in L(u): \\ Z \neq W}} I(z)I(w) = g_0[u] \tag{1}$$

*where $L(u)$ is the set of leaves below $u$.*

*Proof.* Let $\mathcal{S}(u)$ be the set of singletons below vertex $u$, and let $\mathcal{A}(u)$ be the set of artificial taxa below $u$. Then,

$$\sum_{\substack{z,w \in L(u): \\ Z \neq W}} I(z)I(w) = \sum_{\substack{z,w \in \mathcal{L}(u): \\ Z,W \in \mathcal{S}(u), Z \neq W}} I(z)I(w) + \sum_{\substack{z,w \in L(u): \\ Z \in \mathcal{S}(u), W \in \mathcal{A}(u)}} I(z)I(w) + \sum_{\substack{z,w \in L(u): \\ Z,W \in \mathcal{A}(u), Z \neq W}} I(z)I(w)$$

$$= \sum_{\substack{z,w \in L(u): \\ Z,W \in \mathcal{S}(u), Z \neq W}} 1$$

$$+ \left( \sum_{\substack{z \in L(u): \\ Z \in \mathcal{S}(u)}} I(z) \right) \cdot \left( \sum_{\substack{w \in L(u): \\ W \in \mathcal{A}(u)}} I(w) \right)$$

$$+ \frac{\left( \sum_{Z \in \mathcal{A}(u)} \sum_{m \in L(u): M=Z} I(m) \right)^2 - \left( \sum_{Z \in \mathcal{A}(u)} (\sum_{m \in L(u): m=Z} I(m))^2 \right)}{2}$$

$$= \binom{c_0[u]}{2} + c_0[u] \cdot G_1[u] + \frac{G_1[u]^2 - G_2[u]}{2}$$

$$= g_0[u]$$

where $G_1[u] = \sum_{D \in \mathcal{A}(u)} c_D[u]$, and $G_2[u] = \sum_{D \in \mathcal{A}(u)} c_D[u]^2$. □

Lastly, we need to compute the good edges $\mathbb{G}[X, Y]$, which is the total weight of quartets in which $X, Y$ are not siblings. This can be done in constant time, following Lemma 2.

**Lemma 2.** *Let $T$ be a multi-labeled gene tree $T$, and let $X, Y$ be singletons. Then,*

$$\mathbb{G}[X, Y] + \mathbb{B}[X, Y] = \binom{c_0[r] - 2}{2} + (c_0[r] - 2) \cdot G_1[r] + \frac{G_1[r]^2 - G_2[r]}{2} \tag{2}$$

*where $r$ is the root of $T$.*

*Proof.*

$$\begin{aligned}\mathbb{G}[X,Y]+\mathbb{B}[X,Y] &= \sum_{\substack{z,w\in L(r):\\ Z\neq W\neq X\neq Y}} I(x)I(y)I(z)I(w)\\ &= \sum_{\substack{z,w\in L(r):\\ Z\neq W\neq X\neq Y}} I(z)I(w)\\ &= \sum_{\substack{z,w\in L(u):\\ Z,W\in\mathcal{S}(u)\setminus\{X,Y\},\\ Z\neq W}} I(z)I(w) + \sum_{\substack{z,w\in L(u):\\ Z\in\mathcal{S}(u)\setminus\{X,Y\},\\ W\in\mathcal{A}(u)}} I(z)I(w) + \sum_{\substack{z,w\in L(u):\\ Z,W\in\mathcal{A}(u),\\ Z\neq W}} I(z)I(w)\\ &= \binom{c_0[r]-2}{2} + (c_0[r]-2)\cdot G_1[r] + \frac{G_1[r]^2 - G_2[r]}{2}\end{aligned}$$

□

#### 5.2 Quartet Graph Construction Case 2 (Singleton and Artificial Taxon)

In this section, we consider the computation of $\mathbb{B}[X,Y]$ when $X$ is a singleton and $Y$ is an artificial taxon (or vice versa). Our previous algorithm computed $\mathbb{B}[X,Y]$ when it reached the lowest common ancestor (LCA) $X$ and $Y$ during a postorder traversal. However, there are now multiple multiple leaves labeled $Y$ and thus there can be multiple LCAs. Alternatively, we might count the number of quartets with $X$ and $Y$ as siblings that have vertex $v$ as their LCA, meaning that $X$ labels a leaf below vertex $v.left$ and $Y$ labels a leaf below vertex $v.right$ or vice versa. We let $\mathbb{B}_v[X,Y]$ denote the first quantity and $\mathbb{B}_v[Y,X]$ denote the second quantity, defining $\mathbb{A}_v[X,Y]$, $\mathbb{L}_v[X,Y]$, and $\mathbb{R}_v[X,Y]$ in the natural way. Now the total number of quartets with $X,Y$ as siblings can be computed as

$$\mathbb{B}[X,Y] = \mathbb{B}[Y,X] = \sum_{v\in V(T)} \big(\mathbb{B}_v[X,Y] + \mathbb{B}_v[Y,X]\big) \tag{3}$$

Without loss of generality, we show how to compute $\mathbb{B}_v[X,Y]$, where $X$ is a singleton and $Y$ is an artificial taxon. Let's assume this quantity is not zero so $X$ labels *exactly one* leaf below $v.left$ and $Y$ labels *at least one* leaf below $v.right$. In our previous algorithm, we utilized an approach for counting number of ways to select two $z,w$ below vertex $u$ such that $Z\neq W$. Now we must additionally require $Z\neq Y$ because $Y$ already labels a leaf in the quartet. This gives us the modified binomial coefficient

$$g_Y[u] = \binom{c_0[u]}{2} + \Big(c_0[u]\cdot(G_1[u] - c_Y[u])\Big) + \left(\frac{(G_1[u]-c_Y[u])^2 - (G_2[u]-c_Y[u]^2)}{2}\right) \tag{4}$$

from which can compute the modified prefix of $v$: $p_Y[v] = p_Y[v.parent] + g_Y[v.sibling]$. Now we compute a vector $g$ of size $b+1$ and a vector $p$ of size $b$ but note that the time complexity of preprocessing phase is still $O(bn)$. Afterward, we can compute the quantities:

$$\mathbb{A}_v[X,Y] = c_Y[v.right]\cdot g_Y[v.above] \tag{5}$$

$$\mathbb{L}_v[X,Y] = c_Y[v.right]\cdot(p_Y[x] - p_Y[v.left]) \tag{6}$$

These are the same formulas given in the appendix, except that we now exclude $Y$ when taking the modified binomial coefficient and then multiply by $c_Y[v.right]$ (because we can select any leaf labeled $Y$ below $v.right$).

Computing $\mathbb{R}_v[X,Y]$ is more complicated, and thus, we define a *different* dynamic programming algorithm that counts triplets of the form $((W,Z),Y)$. Specifically, we define

$$\mathbb{R}_v[X,Y] = f_{Y,0}[v.right] \tag{7}$$

where $f_{Y,0}[u]$ is the number of triplets below vertex $u = v.right$ of the form $((W,Z),Y)$. This quantity can be computed in $O(n)$ time a preorder traversal, after initializing $f_{y,0}[leaf] = 0$, as follows:

$$f_{Y,0}[u] = f_{Y,0}[u.left] + f_{Y,0}[u.right] + \Big(c_Y[u.left]\cdot g_Y[u.right]\Big) + \Big(c_Y[u.right]\cdot g_Y[u.left]\Big) \tag{8}$$

where the first and second term come from selecting three leaves below $u.left$ and $u.right$, respectively (these triplets have already been counted). The last two terms come from counting triplets whose LCA is $u$. We repeat this preprocessing to compute $f$ for all $b$ artificial taxa; thus, the total time complexity of the preprocessing phase is still $O(bn)$.

After the preprocessing phase, we can compute $\mathbb{B}_v[X, Y]$ in constant time (a similar argument can be made for $\mathbb{B}_v[Y, X]$). Both of these quantities will equal zero (and do not need to be computed) for vertices that are not on the path from the singleton $X$ to the root; thus, we can compute $\mathbb{B}[X, Y]$ in $O(n)$ time. Repeating this calculation for all ways of selecting one singleton and one artificial taxon gives us the final time complexity: $O(abh + bn)$.

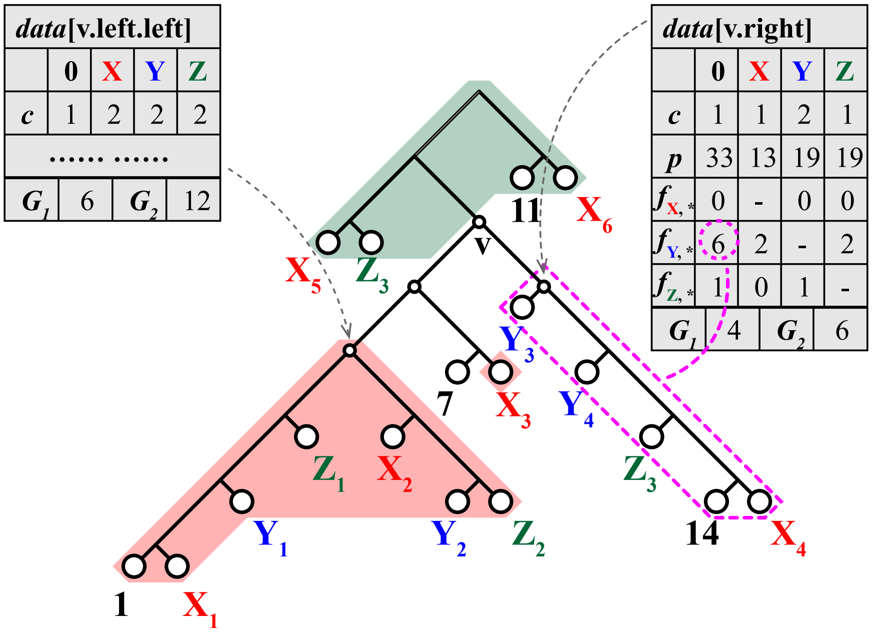

Figure S13: $\mathbb{B}_v$ **with one artificial taxon.** Above is an example of the data stored to compute $\mathbb{B}_v[7, Y]$, where 7 is a singleton in $v.left$ and $Y$ is an artificial taxa in $v.right$. $\mathbb{B}_v[7, Y] = \mathbb{A}_v[7, Y] + \mathbb{L}_v[7, Y] + \mathbb{R}_v[7, Y] = 2 \cdot 5 + 2 \cdot 8 + 6 = 32$.

#### 5.3 Quartet Graph Construction Case 3 (Two Artificial Taxa)

Lastly, we consider the computation of $\mathbb{B}[X, Y]$ when $X$ and $Y$ are both artificial taxa. This largely proceeds as previously described in the previous Section. First, we update the binomial coefficient to exclude both $X$ and $Y$ (instead of just $Y$) as follows:

$$g_{X,Y}[v] = \binom{c_0[v]}{2} + \Big(c_0[v] \cdot (G_1[v] - c_Y[v] - c_X[v])\Big) + \left(\frac{(G_1[v] - c_Y[v] - c_X[v])^2 - (G_2[v] - c_Y[v]^2 - c_X[v]^2)}{2}\right)$$

Second, we update the quantity for counting triplets

$$f_{Y,X}[v] = f_{Y,X}[v.left] + f_{Y,X}[v.right] + c_Y[v.left] \cdot g_{X,Y}[v.right] + c_Y[v.right] \cdot g_{X,Y}[v.left] \quad (9)$$

Third, we update quantities associated with $\mathbb{B}_v[X, Y]$, specifically

$$\mathbb{A}_v[X, Y] = c_X[v.left] \cdot c_Y[v.right] \cdot g_{X,Y}[v.above] \quad (10)$$
$$\mathbb{L}_v[X, Y] = c_Y[v.right] \cdot f_{X,Y}[v.left] \quad (11)$$
$$\mathbb{R}_v[X, Y] = c_X[v.left] \cdot f_{Y,X}[v.right] \quad (12)$$

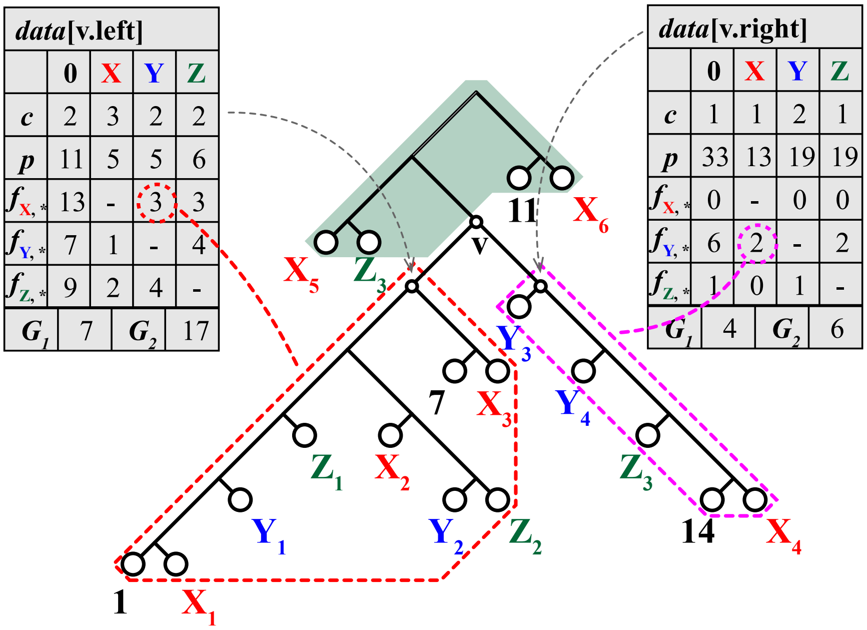

Figure S14: $\mathbb{B}_v$ **with two artificial taxa.** In subfigure b), we show how to compute $\mathbb{B}_v[X, Y]$ where both $X$ and $Y$ are artificial. Again, $\mathbb{B}_v[X, Y]$ is given by the sum $\mathbb{A}_v[X, Y]$, $\mathbb{L}_v[X, Y]$, and $\mathbb{R}_v[X, Y]$. $\mathbb{A}_v[X, Y]$ is still derived from the green subtree above $v$ except that not only $Y$ but also $X$ is excluded. Consequently, $g_{X,Y}[v.above] = 1$ since there are only two taxa in the subtree except $X$, and $\mathbb{A}_v[X, Y]$ is therefore $c_X[v.right] \cdot c_Y[v.right] \cdot g_{X,Y}[v.above] = 6$. On the other hand, $\mathbb{L}_v[X, Y]$ is $c_Y[v.right] \cdot f_{X,Y}[v.left] = 2 \times 3 = 6$ while $\mathbb{R}_v[X, Y] = 3 \times 2 = 6$ because $c_X[v.left] = 3$ and $f_{Y,X}[v.right] = 2$. Finally, we have $\mathbb{B}_v[X, Y] = 6 + 6 + 6 = 18$. Similarly, $\mathbb{B}_v[Y, X] = 2 + 1 + 0 = 3$ because $c_Y[v.left] = 2$, $c_X[v.right] = 1$, $g_{X,Y}[v.above] = 1$, $f_{X,Y}[v.right] = 0$ and $f_{Y,X}[v.left] = 1$.

The key thing to note is that we are now storing a matrix of size $b^2$ at each vertex $v$. This brings the preprocessing phase to $O(b^2 n)$, after which we can compute $\mathbb{B}_v[X, Y]$ in constant time. This quantity could be non-zero for any internal vertex of $T$ so the total time to compute $\mathbb{B}[X, Y]$ is once again $O(n)$. We repeat this calculation for all ways of selecting one two artificial taxa to get the total time complexity is $O(b^2 n)$.

#### 5.4 Algorithms

**Algorithm 1:** Initialization

**Data:** the input gene tree $T$ and pre-computed $c[v]$ and $g[v]$ values

**Result:** the $p[v]$ and $f[v]$ values for each node $v$ in $T$

**for** *each node $v$ of $T$ in the pre-order* **do**
    **if** *$v$ is the root* **then**
        $p_0[v] \leftarrow 0$
        **for** *each artificial taxa $X$* **do**
            $p_X[v] \leftarrow 0$
    **else**
        **if** *$v$ is not a leaf* **then**
            $p_0[v] \leftarrow p_0[v.parent] + g_0[v.sibling]$
            **for** *each artificial taxa $X$* **do**
                $p_X[v] \leftarrow p_X[v.parent] + g_X[v.sibling]$

**for** *each node $v$ of $T$ in the post-order* **do**
    **if** *$v$ is a leaf* **then**
        **for** *each artificial taxa $Y$* **do**
            $f_{Y,0}[v] \leftarrow 0$
        **for** *each artificial taxa $Y$* **do**
            **for** *each artificial taxa $X \neq Y$* **do**
                $f_{Y,X}[v] \leftarrow 0$
    **else**
        **for** *each artificial taxa $Y$* **do**
            $f_{Y,0}[v] \leftarrow f_{Y,0}[v.left] + f_{Y,0}[v.right] \ + c_Y[v.left] \cdot g_Y[v.right] + c_Y[v.right] \cdot g_Y[v.left]$
        **for** *each artificial taxa $Y$* **do**
            **for** *each artificial taxa $X \neq Y$* **do**
                $f_{Y,X}[v] \leftarrow$
                  $f_{Y,X}[v.left] + f_{Y,X}[v.right] \ + c_Y[v.left] \cdot g_{X,Y}[v.right] + c_Y[v.right] \cdot g_{X,Y}[v.left]$

**Algorithm 2:** Quartet Graph Construction – Bad Edges

**Data:** the input gene tree $T$ with all pre-computed values
**Result:** $\mathbb{B}[X,Y]$ : the weights of the bad edges between each pair of $X$ and $Y$

**for** *each internal node* $v$ *in* $T$ **do**
  **for** *each taxa* $X$ *below* $v.left$ **do**
    **for** *each taxa* $Y$ *below* $v.right$ *such that* $X \neq Y$ **do**
      **if** $X$ *is an artificial taxa* **then**
        **if** $Y$ *is an artificial taxa* **then**
          $\mathbb{A}_v[X,Y] \leftarrow c_X[v.left] \cdot c_Y[v.right] \cdot g_{X,Y}[v.above]$
          $\mathbb{L}_v[X,Y] \leftarrow c_Y[v.right] \cdot f_{X,Y}[v.left]$
          $\mathbb{R}_v[X,Y] \leftarrow c_X[v.left] \cdot f_{Y,X}[v.right]$
        **else**
          $\mathbb{A}_v[X,Y] \leftarrow c_Y[v.right] \cdot g_Y[v.above]$
          $\mathbb{L}_v[X,Y] \leftarrow c_Y[v.right] \cdot (p_Y[x] - p_Y[v.left])$ where $y$ is the unique leaf labeled $Y$
          $\mathbb{R}_v[X,Y] \leftarrow f_{Y,0}[v.right]$
      **else**
        **if** $Y$ *is an artificial taxa* **then**
          $\mathbb{A}_v[X,Y] \leftarrow c_X[v.left] \cdot g_X[v.above]$
          $\mathbb{L}_v[X,Y] \leftarrow f_{X,0}[v.left]$
          $\mathbb{R}_v[X,Y] \leftarrow c_X[v.left] \cdot (p_X[y] - p_X[v.right])$
        **else**
          $\mathbb{A}_v[X,Y] \leftarrow g_0[v.above]$
          $\mathbb{L}_v[X,Y] \leftarrow p_0[x] - p_0[v.left]$
          $\mathbb{R}_v[X,Y] \leftarrow p_0[y] - p_0[v.right]$
      $\mathbb{B}[X,Y] \leftarrow \mathbb{B}[X,Y] + \mathbb{A}_v[X,Y] + \mathbb{L}_v[X,Y] + \mathbb{R}_v[X,Y]$
      $\mathbb{B}[X,Y] \leftarrow \mathbb{B}[X,Y] + \mathbb{A}_v[X,Y] + \mathbb{L}_v[X,Y] + \mathbb{R}_v[X,Y]$

**Algorithm 3:** Quartet Graph Construction – Good Edges

**Data:** the input gene tree $T$ with root $r$, all pre-computed values for $T$, and $\mathbb{B}$
**Result:** $\mathbb{G}[X,Y]$ : the weights of the good edges between each pair of $X$ and $Y$

**for** *each taxa* $X$ **do**
  **for** *each taxa* $Y$ **do**
    **if** $X$ *is an artificial taxa* **then**
      **if** $Y$ *is an artificial taxa* **then**
        $\mathbb{G}[X,Y] = g_{X,Y}[r] \cdot c_X[r] \cdot c_Y[r] - \mathbb{B}[X,Y]$
      **else**
        $\mathbb{G}[X,Y] = \left(\binom{c_0[r]-1}{2} + \left((c_0[r]-1) \cdot (G_1[v] - c_X[v])\right) + \left(\frac{(G_1[r]-c_X[r])^2-(G_2[r]-c_X[r]^2)}{2}\right)\right) \cdot c_X[r] - \mathbb{B}[X,Y]$
    **else**
      **if** $Y$ *is an artificial taxa* **then**
        $\mathbb{G}[X,Y] = \left(\binom{c_0[r]-1}{2} + \left((c_0[r]-1) \cdot (G_1[r] - c_Y[r])\right) + \left(\frac{(G_1[r]-c_Y[r])^2-(G_2[r]-c_Y[r]^2)}{2}\right)\right) \cdot c_Y[r] - \mathbb{B}[X,Y]$
      **else**
        $\mathbb{G}[X,Y] = \binom{c_0[r]-2}{2} + (c_0[r]-2) \cdot G_1[r] + \frac{G_1[r]^2-G_2[r]}{2} - \mathbb{B}[X,Y]$

##### 5.5 Time Complexity

**Theorem 1.** *Let $n$ leaves in the gene tree $T$, and let $s$ be the number of leaf labels for the current subproblem. Then, we compute $\mathbb{B}$ and $\mathbb{G}$ in $O(s^2 n)$ time using Algorithms 1–3.*

*Proof.* Let $a$ be the number of singletons in the subproblem, and let $b$ be the number of artificial taxa (so $a + b = s$). To compute $\mathbb{B}$, we need to compute $\mathbb{A}, \mathbb{L}, \mathbb{R}$, which in turn depend on other quantities that are computed during a preprocessing phase.

**Preprocessing phase:** Computing the **number of taxa** below vertex $v$:

- $c_0[v]$ takes $O(n)$ time
- $c_Y[v]$ for each artificial taxon $Y$ takes $O(bn)$ time
- all $G_1[v]$ and $G_2[v]$ takes $O(bn)$ time

via a post-order traversal on $T$. Computing the **modified binomial coefficients**:

- all $g_0[v]$ takes $O(n)$ time
- all $g_Y[v]$ for each artificial taxon $Y$ takes $O(bn)$ time
- all $g_{X,Y}[v]$ for each pair $X, Y$ of artificial taxa takes $O(b^2 n)$ time

using $G_1[v]$ and $G_2[v]$. Computing the **prefixes**

- $p_0[v]$ takes $O(n)$ time
- $p_X[v]$ for each artificial taxon $X$ takes $O(bn)$ time

see lines 1-10 of Algorithm 1. Computing the **triplet counters**

- $f_{Y,0}[v]$ for each artificial taxon $Y$ takes $O(bn)$ time
- $f_{Y,X}[v]$ for each pair $X, Y$ of artificial taxa takes $O(b^2 n)$ time

see lines 11-23 of Algorithm 1. To sum up, the total initialization time is $O(b^2 n)$ for each gene tree.

Now we can compute the weights of the **bad edges** $\mathbb{B}$ via Algorithm 2. Each iteration of Algorithm 2 (lines 4-23) costs $O(1)$ time. In other words, given any triple of $v$, $X$, and $Y$, the corresponding $\mathbb{B}_v[X, Y]$ is computed in constant time. Consequently, the total time complexity depends on the number of the distinct tripples $(v, X, Y)$. If both $X$ and $Y$ are singletons, then there is only one unique $v$ at which $\mathbb{B}[X, Y]$ or $\mathbb{B}[Y, X]$ are updated because $v$ must be the LCA of $X$ and $Y$. As a result, the triplet of $(v, X, Y)$ is unique when $X$ and $Y$ are singleton, and the number of triplets for all singleton pairs of $X$ and $Y$ is $O(a^2)$. If $X$ is a singleton and $Y$ is an artificial taxon, the number of $v$ is $O(h)$, where $h$ is the height of $T$, because $v$ must be one of the ancestors of $X$. In this case, the number of the triplets is $O(abh)$. The same holds in the case where $Y$ is a singleton and $X$ is an artificial taxa. If both $X$ and $Y$ are artificial taxa, then the number of $v$ is $O(n)$ since $v$ could be any inner node in the tree, and hence the number of triplets is $O(b^2 n)$. To sum up, the total number of $(v, X, Y)$, as well as the time complexity of Algorithm 2, is $O(a^2 + abh + b^2 n) = O(s^2 n)$. In the last step, we compute the weights of the **good edges** $\mathbb{G}$. We can update each $\mathbb{G}$ in $O(s^2)$ time using Algorithm 3. Thus, the time complexity is $O(s^2)$.

To conclude, the total time complexity is $O(s^2 n)$. □

**Theorem 2.** *Let $n$ be the number of species and let $k$ be the number of gene trees. TREE-QMC has time complexity $O(n^3 k)$ if subproblems are produced in a perfectly balanced fashion.*

*Proof.* At each step in the divide phase of the algorithm, we compute a bipartition and then recurse on two subproblems. Each subproblem is defined by taking the taxa on one side of the bipartition and adding an artificial taxon to represent the taxa on the other side of the bipartition. Thus, a subproblem of size $2^p + 2$ is divided into two subproblems of size $\frac{2^p+2}{2} + 1 = 2^{p-1} + 2$ if the division is balanced. The work $f$ per subproblem has time complexity $O(s^3 + s^2 nk)$ where the first term comes from applying the heuristic for max cut and the second term comes from constructing the quartet graph by applying our algorithm to all gene trees (Theorem 1). Therefore,

$$
\begin{aligned}
T(2^p+2) &= \sum_{i=0}^{p} 2^i \cdot f(2^{p-i}+2) \quad \text{where } f \text{ is } O(s^3+s^2nk) \\
&= \sum_{i=0}^{p} 2^i \cdot [(2^{p-i}+2)^3 + (2^{p-i}+2)^2 \cdot (2^p+2) \cdot k] \\
&= \sum_{i=0}^{p} 2^i \cdot (2^{p-i}+2)^3 + \sum_{i=0}^{p} 2^i \cdot (2^{p-i}+2)^2 \cdot (2^p+2) \cdot k \\
&= \sum_{i=0}^{p} 2^i \cdot (2^{p-i}+2)^3 + (2^p+2) \cdot k \cdot \sum_{i=0}^{p} 2^i \cdot (2^{p-i}+2)^2 \\
&= \sum_{i=0}^{p} 2^i \cdot (2^{3(p-i)} + 3 \cdot 2^{2(p-i)+1} + 3 \cdot 2^{p-i+2} + 2^3) + (2^p+2) \cdot k \cdot \sum_{i=0}^{p} 2^i \cdot (2^{2(p-i)} + 2^{p-i+2} + 2^2) \\
&= \sum_{i=0}^{p} (2^{3(p-i)+i} + 3 \cdot 2^{2(p-i)+1+i} + 3 \cdot 2^{p-i+2+i} + 2^{3+i}) + (2^p+2) \cdot k \cdot \sum_{i=0}^{p} (2^{2(p-i)+i} + 2^{p-i+2+i} + 2^{2+i}) \\
&= \Big(\frac{1}{3} \cdot 2^p \cdot (4^p-1) + 3 \cdot 2^{p+1} \cdot (2^{p+1}-1) + 12 \cdot (2^{p+1}-1) + 2^{p+4} - 8\Big) \\
&\quad + (2^p+2) \cdot k \cdot \big(2^p \cdot (2^{p+1}-1) + 2^{p+2} \cdot (p+1) + 2^{p+3} - 4\big) \\
&= O(2^{3p} + 2^p \cdot k \cdot 2^{2p}) \\
&= O(k \cdot 2^{3p}) \\
&= O(kn^3)
\end{aligned}
$$

□

##### 5.6 Correctness

**Lemma 3.** *Let $v$ be an internal node in $T$, let $X$ be a taxon below $v.left$, and let $Y$, $Z$, and $W$ be three taxa below $v.right$. The total weight of quartets $\{X, Y, Z, W\}$ in which $X$ and $Y$ are siblings, denoted by $\mathbb{R}_v[X, Y]$, is*

$$
\begin{cases}
c_X[v.left] \cdot f_{Y,X}[v.right] & \textit{if both } X \textit{ and } Y \textit{ are artificial} \\
f_{Y,0}[v.right] & \textit{if only } Y \textit{ is artificial} \\
c_x[v.left] \cdot (p_X[y] - p_x[v.right]) & \textit{if only } X \textit{ is artificial} \\
p_0[y] - p_0[v.right] & \textit{if both } X \textit{ and } Y \textit{ are singletons}
\end{cases}
\tag{13}
$$

*where the values of $p_0[v]$, $p_X[v]$, $f_{Y,0}[v]$, and $f_{Y,X}[v]$ are given by Algorithm 1. Similarly, when $Z$ and $W$ are from $v.left$, the total weight, denoted by $\mathbb{L}_v[X, Y]$, is*

$$
\begin{cases}
c_y[v.right] \cdot f_{X,Y}[v.left] & \textit{if both } X \textit{ and } Y \textit{ are artificial} \\
f_{X,0}[v.left] & \textit{if only } Y \textit{ is artificial} \\
c_Y[v.right] \cdot (p_Y[x] - p_Y[v.left]) & \textit{if only } X \textit{ is artificial} \\
p_0[x] - p_0[v.left] & \textit{if both } X \textit{ and } Y \textit{ are singletons}
\end{cases}
\tag{14}
$$

*Finally, when $Z$ and $W$ are from $v.above$, the total weight, denoted by $\mathbb{A}_v[X, Y]$, is*

$$
\begin{cases}
c_X[v.left] \cdot c_Y[v.right] \cdot g_{X,Y}[v.above] & \textit{if both } X \textit{ and } Y \textit{ are artificial} \\
c_Y[v.left] \cdot g_Y[v.above] & \textit{if only } Y \textit{ is artificial} \\
c_X[v.right] \cdot g_X[v.above] & \textit{if only } X \textit{ is artificial} \\
g_0[v.above] & \textit{if both } X \textit{ and } Y \textit{ are singletons}
\end{cases}
\tag{15}
$$

*Proof.* Note: capital letters are used to denote the label of a leaf. For example, leaf $m$ would have label $M$ in the current subproblem.

The proof for $\mathbb{A}_v[X,Y]$ is simple since there is only one subtree above $v$. We prove the case in which both $X$ and $X$ are artificial,

$$\begin{aligned}\mathbb{A}_v[X,Y] &= \sum_{\substack{m\in L(v.left):\\ M=X}} \sum_{\substack{n\in L(v.right)\\ N=Y}} \sum_{\substack{z,w\in L(v.above):\\ Z\neq Y\neq X\neq W}} I(m)I(n)I(z)I(w)\\ &= \sum_{\substack{m\in L(v.left):\\ M=X}} I(m)\cdot\Bigg(\sum_{\substack{n\in L(v.right):\\ N=Y}} I(n)\cdot\Bigg(\sum_{\substack{z,w\in L(v.above):\\ Z\neq W\neq X\neq Y}} I(z)I(w)\Bigg)\Bigg)\\ &= c_X[v.left]\cdot c_Y[v.right]\cdot g_{X,Y}[v.above],\end{aligned}$$

These equations are for artificial taxa $X, Y$ but we can allow $X$ to be a singleton by setting $c_X[u] = I(x) = 1$ and decreasing $c_0[u]$ by $I(x) = 1$ when computing $g_{X,Y}[u]$ (and similarly to allow $Y$ to be a singleton). The other three cases of $\mathbb{A}_v[X,Y]$ are similar. In the following, we focus on $\mathbb{R}_v[X,Y]$ because the proof for $\mathbb{L}_v[x,y]$ is the same due to the symmetry. Recall that $X$ is a taxon below $v.left$, and $Y$, $Z$, and $W$ be three taxa below $v.right$.

**If both $X$ and $Y$ are singletons**, the prefix-sum technique is applied:

$$\begin{aligned}\mathbb{R}_v[X,Y] &= \sum_{\substack{z,w\in L(v.right):\\ Z\neq W\neq Y\\ q(x,y,z,w)=x,y|z,w}} I(x)I(y)I(z)I(w)\\ &= \sum_{\substack{z,w\in L(v.right):\\ Z\neq W\neq Y\\ q(x,y,z,w)=x,y|z,w}} I(z)I(w)\\ &= \sum_{u\in\mathcal{P}(v.right,y)} \sum_{\substack{z,w\in L(u)\\ Z\neq W}} I(z)I(w)\\ &= \sum_{u\in\mathcal{P}(v.right,y)} g_0[u]\\ &= \sum_{u\in\mathcal{P}(root,y)} g_0[u] - \sum_{u\in\mathcal{P}(root,v.right)} g_0[u]\\ &= p_0[y] - p_0[v.right],\end{aligned}$$

where $\mathcal{P}(i,j)$ is the set of the subtrees adjacent to the path from leaves $i$ (included) to $j$ (excluded). Note that $g_0[u]$ is used to compute $\sum_{z,w\in\mathcal{L}(u),Z\neq W} I(z)I(w)$ according to Lemma 1 in the main text.

**If $X$ is a singleton while $Y$ is an artificial taxon**, there could be multiple leaves labeled by $Y$ in $L(v.right)$. Consequently, $Y$ must be excluded when $Z$ and $W$ are selected, and thus $g_Y[u]$ is used instead of $g_0[u]$:

$$\begin{aligned}\mathbb{R}_v[X,Y] &= \sum_{\substack{n\in L(v.right):\\ N=Y}} \sum_{\substack{z,w\in L(v.right):\\ Z\neq W\neq Y\\ q(x,n,z,w)=x,n|z,w}} I(x)I(n)I(z)I(w)\\ &= \sum_{\substack{n\in L(v.right):\\ N=Y}} I(n)\cdot\Bigg(\sum_{u\in\mathcal{P}(v.right,n)} \sum_{\substack{z,w\in L(u):\\ Z\neq W\neq Y}} I(z)I(w)\Bigg)\\ &= \sum_{\substack{n\in L(v.right):\\ N=Y}} I(n)\cdot \sum_{u\in\mathcal{P}(v.right,n)} g_Y[u]\\ &= f_{Y,0}[v.right]\end{aligned}$$

Algorithm 1 computes $f_{Y,0}[v]$ according to the following recurrence:

$$f_{Y,0}[v] = \begin{cases} 0 & \text{if } v \text{ is a leaf} \\ f_{Y,0}[v.left] + f_{Y,0}[v.right] & \\ +c_Y[v.left] \cdot g_Y[v.right] + c_Y[v.right] \cdot g_Y[v.left] & \text{if } v \text{ is an inner node} \end{cases}.$$

The correctness of the recurrence is shown as follows.

$$\begin{aligned}
f_{Y,0}[v] &= \sum_{\substack{n \in L(v): \\ N=Y}} I(n) \cdot \sum_{u \in \mathcal{P}(v,n)} g_Y[u] \\
&= \sum_{\substack{n \in L(v.left): \\ N=Y}} I(n) \sum_{u \in \mathcal{P}(v,n)} g_y[u] + \sum_{\substack{n \in L(v.right): \\ N=Y}} I(n) \sum_{u \in \mathcal{P}(v,n)} g_Y[u] \\
&= \sum_{\substack{n \in L(v.left): \\ N=Y}} I(n) \cdot \left( g_Y[v.right] + \sum_{u \in \mathcal{P}(v.left,n)} g_y[u] \right) \\
&\quad + \sum_{\substack{n \in L(v.right): \\ N=Y}} I(n) \cdot \left( g_Y[v.left] + \sum_{u \in \mathcal{P}(v.right,n)} g_Y[u] \right) \\
&= \sum_{\substack{n \in L(v.left): \\ N=Y}} I(n) \cdot \sum_{u \in \mathcal{P}(v.left,n)} g_Y[u] + \sum_{\substack{n \in L(v.right): \\ N=Y}} I(n) \cdot \sum_{u \in \mathcal{P}(v.right,n)} g_Y[u] \\
&\quad + g_Y[v.right] \cdot \sum_{\substack{n \in L(v.left): \\ N=Y}} I(n) + g_Y[v.left] \cdot \sum_{\substack{n \in L(v.right): \\ N=Y}} I(n) \\
&= f_{Y,0}[v.left] + f_{Y,0}[v.right] + c_Y[v.left] \cdot g_Y[v.right] + c_Y[v.right] \cdot g_y[v.left].
\end{aligned}$$

**If $X$ is an artificial taxon and $Y$ is a singleton**, $X$ could also appear in $L(v.right)$ so we must be excluded to select $Z$ and $W$, and therefore $g_X[u]$ is used to compute the sum of $I(z)I(w)$ in a subtree.

$$\begin{aligned}
\mathbb{R}_v[x,y] &= \sum_{\substack{m \in L(v.left) \\ M=X}} \sum_{\substack{z,w \in L(v.right): \\ Z \neq W \neq X \\ q(m,y,z,w)=m,y|z,w}} I(m)I(y)I(z)I(w) \\
&= \sum_{\substack{m \in L(v.left): \\ M=X}} I(m) \sum_{u \in \mathcal{P}(v.right,y)} \sum_{\substack{z,w \in L(u): \\ Z \neq W \neq X}} I(z)I(w) \\
&= \sum_{\substack{m \in L(v.left): \\ M=X}} I(m) \cdot \left( \sum_{u \in \mathcal{P}(v.right,y)} \sum_{\substack{z,w \in L(u): \\ Z \neq W \neq X}} I(z)I(w) \right) \\
&= \sum_{\substack{m \in L(v.left): \\ M=X}} I(m) \cdot \left( \sum_{u \in \mathcal{P}(v.right,y)} g_x[u] \right) \\
&= \sum_{\substack{m \in L(v.left): \\ M=X}} I(m) \cdot \left( \sum_{u \in \mathcal{P}(root,y)} g_X[u] - \sum_{u \in \mathcal{P}(root,v.right)} g_X[u] \right) \\
&= \sum_{\substack{m \in L(v.left): \\ M=X}} I(m) \cdot (p_X[y] - p_X[v.right]) \\
&= c_X[v.left] \cdot (p_X[y] - p_X[v.right]).
\end{aligned}$$

**Finally, if both $X$ and $Y$ are artificial taxa, $g_{X,Y}[u]$ is used.**

$$
\begin{aligned}
\mathbb{R}_v[X,Y] =& \sum_{\substack{m\in L(v.left):\\ M=X}} \sum_{\substack{N\in L(v.right):\\ n=Y}} \sum_{\substack{z,w\in L(v.right):\\ Z\neq W\neq X\neq Y\\ q(m,n,z,w)=m,n|z,w}} I(m)I(n)I(z)I(w)\\
=& \sum_{\substack{m\in L(v.left):\\ M=X}} I(m)\cdot\Bigg(\sum_{\substack{n\in L(v.right):\\ N=Y}} I(n)\cdot\Bigg(\sum_{u\in\mathcal{P}(v.right,n)} \sum_{\substack{z,w\in\mathcal{L}(u):\\ Z\neq W\neq X\neq Y}} I(z)I(w)\Bigg)\Bigg)\\
=& \sum_{\substack{m\in L(v.left):\\ M=X}} I(m)\cdot\Bigg(\sum_{\substack{n\in L(v.right):\\ N=Y}} I(n)\cdot\Bigg(\sum_{u\in\mathcal{P}(v.right,n)} g_{X,Y}[u]\Bigg)\Bigg)\\
=& \sum_{\substack{m\in L(v.left):\\ M=X}} I(m)\cdot f_{Y,X}[v.left]\\
=& c_X[v.left]\cdot f_{Y,X}[v.left]
\end{aligned}
$$

Algorithm 1 computes $f_{Y,X}[v]$ according to the following recurrence:

$$
f_{Y,X}[v] = \begin{cases} 0 & \text{if } v \text{ is a leaf}\\ f_{Y,X}[v.left] + f_{Y,X}[v.right] & \\ +c_Y[v.left]\cdot g_{X,Y}[v.right] + c_Y[v.right]\cdot g_{X,Y}[v.left] & \text{if } v \text{ is an inner node} \end{cases}
$$

whose proof of correctness is similar to $f_{Y,0}[v]$, except that $g_{X,Y}[u]$ is used instead of $g_X[u]$. □

**Lemma 4.** *Let $T$ be a binary tree, and let $X$ and $Y$ be taxa ($X \neq Y$). Then,*

$$
\mathbb{B}[X,Y] = \sum_{v\in V(T)} \Big(\mathbb{A}_v[X,Y] + \mathbb{L}_v[X,Y] + \mathbb{R}_v[X,Y]\Big) + \Big(\mathbb{A}_v[Y,X] + \mathbb{L}_v[Y,X] + \mathbb{R}_v[Y,X]\Big)
$$

*where $V(T)$ is the set of internal nodes in gene tree $T$.*

*Proof.* By definition, $\mathbb{B}[X,Y]$ is the total weight of the quartet in $T$ whose topology is $X,Y|Z,W$, where $Z$

and $W$ are two other taxa, so we have

$$\begin{aligned}
\mathbb{B}[X,Y] &= \sum_{\substack{m\in L(T):\\ M=X}} \sum_{\substack{n\in L(T):\\ N=Y}} \sum_{\substack{z,w\in L(T):\\ Z\neq W\neq X\neq Y\\ q(n,m,z,w)=m,n|z,w}} I(m,n,z,w) \\
&= \sum_{v\in V(T)} \sum_{\substack{m\in L(v.left):\\ M=X}} \sum_{\substack{n\in L:(v.right)\\ N=Y}} \sum_{\substack{z,w\in L(T):\\ Z\neq W\neq X\neq Y\\ q(m,n,z,w)=m,n|z,w}} I(m,n,z,w) \\
&\quad + \sum_{v\in V(T)} \sum_{\substack{m\in L(v.right):\\ M=X}} \sum_{\substack{n\in L:(v.left)\\ N=Y}} \sum_{\substack{z,w\in \mathcal{L}(T):\\ Z\neq W\neq X\neq Y\\ q(m,n,z,w)=m,n|z,w}} I(m,n,z,w) \\
&= \sum_{v\in V(T)} \sum_{\substack{m\in L(v.left):\\ M=X}} \sum_{\substack{n\in L(v.right):\\ N=Y}} \Bigg( \sum_{\substack{z,w\in L(v.above):\\ Z\neq W\neq X\neq Y\\ q(n,m,z,w)=m,n|z,w}} I(m,n,z,w) \\
&\quad + \sum_{\substack{z,w\in L(v.left):\\ Z\neq W\neq X\neq Y\\ q(m,n,z,w)=m,n|z,w}} I(m,n,z,w) + \sum_{\substack{z,w\in L(v.right):\\ Z\neq W\neq X\neq Y\\ q(m,n,z,w)=m,n|z,w}} I(m,n,z,w)\Bigg) \\
&+ \sum_{v\in V(T)} \sum_{\substack{m\in L(v.right):\\ M=X}} \sum_{\substack{n\in L(v.left):\\ N=Y}} \Bigg( \sum_{\substack{z,w\in L(v.above):\\ Z\neq W\neq X\neq Y\\ q(m,n,z,w)=m,n|z,w}} I(m,n,z,w) \\
&\quad + \sum_{\substack{z,w\in L(v.left):\\ Z\neq W\neq X\neq Y\\ q(m,n,z,w)=m,n|z,w}} I(m,n,z,w) + \sum_{\substack{z,w\in L(v.right):\\ Z\neq W\neq X\neq Y\\ q(m,n,z,w)=m,n|z,w}} I(m,n,z,w)\Bigg)
\end{aligned}$$

According to Lemma 3, this is

$$\sum_{v\in V(T)} \Big(\mathbb{A}_v[X,Y] + \mathbb{L}_v[X,Y] + \mathbb{R}_v[X,Y]\Big) + \Big(\mathbb{A}_v[Y,X] + \mathbb{L}_v[Y,X] + \mathbb{R}_v[Y,X]\Big)$$

□

The following theorem follows from the Lemma above.

**Theorem 3.** *Let $T$ be a binary tree, and let $X, Y$ be taxa ($X \neq Y$). Algorithm 2 computes $\mathbb{B}[X,Y]$ correctly for $T$.*

**Lemma 5.** *Let $X$ and $Y$ be taxa. Then,*

$$\mathbb{G}[X,Y] + \mathbb{B}[X,Y] = g_{X,Y}[r]\cdot c_X[r]\cdot c_Y[r] \tag{16}$$

*where $r$ is the root of gene tree $T$. These equations are for artificial taxa; however, they can be modified to accommodate singletons as follows (see Lemma 2 for details). If $X$ is a singleton, set $c_X[r] = 1$ in the equation above. Then, compute $g_{X,Y}[r]$ by setting $c_X[r]$ to 0 and subtracting 1 from $c_0[r]$. If $Y$ is a singleton, set $c_Y[r] = 1$ in the equation above. Then, compute $g_{X,Y}[r]$ by setting $c_Y[r]$ to 0 and subtracting 1 from $c_0[r]$.*

*Proof.*

$$\begin{aligned}
\mathbb{G}[X,Y]+\mathbb{B}[X,Y] =& \sum_{\substack{m,n,z,w\in L(r): \\ M=X,N=Y, \\ Z\neq W\neq X\neq Y}} I(m,n,z,w) \\
=& \sum_{\substack{m\in L(T): \\ M=X}} I(m)\cdot\Bigg(\sum_{\substack{n\in L(T): \\ N=Y}} I(n)\cdot\Bigg(\sum_{\substack{z,w\in L(T): \\ Z\neq W\neq X\neq Y}} I(z)I(w)\Bigg)\Bigg) \\
=& c_X[T]\cdot c_Y[T]\cdot g_{X,Y}[T].
\end{aligned}$$

Note: capital letters are used to denote the label of a leaf. For example, leaf $m$ would have label $M$ in the current subproblem. □

The theorem below follows from the Theorem and Lemma above.

**Theorem 4.** *Let $T$ be a binary tree, and let $X,Y$ be taxa ($X\neq Y$). Algorithm 3 computes $\mathbb{G}[X,Y]$ correctly for $T$.*
